# Identification of Cholangiocarcinoma (CCA) Subtype-Specific Biomarkers

**DOI:** 10.1101/2023.08.21.554136

**Authors:** Jacob Croft, Liyuan Gao, Odalys Quintanar, Victor Sheng, Jun Zhang

## Abstract

Liver cancer ranks sixth globally in diagnoses and second in cancer-related deaths. Cholangiocarcinoma (CCA), a relatively rare cancer originating from bile duct epithelium, constitutes 2% of all cancers, with increasing occurrences in Westerns. Incidence is influenced by inflammation, genetics, risk factors, and regional disparities, with higher rates in the Eastern hemisphere. Diagnostic and prognostic biomarkers are pivotal for effective cancer prevention and management. Recent research explores serum proteins for non-invasive CCA diagnosis and proposes targeted receptor approaches for therapeutics.

This study aims to identify these biomarkers via bioinformatics analysis of public datasets, focusing on CCA patient transcriptomes to uncover gene biomarkers linked to age and survival. Pathway analysis reveals functions and pathways associated with these biomarkers. Additionally, the study employs the ESM-TFpredict machine learning model to predict transcription factors (TFs) using protein sequence data.

Leveraging publicly available data enhances our understanding of liver cancer’s molecular profiles and clinical relevance, particularly concerning CCA, This study integrates bioinformatics analysis, transcriptomic exploration, and machine learning to unveil a novel set of potential diagnostic and prognostic biomarkers for CCA.

## Introduction

Liver cancer, specifically hepatocellular carcinomas (HCC) and cholangiocarcinomas (CCA), poses a significant global health challenge as the sixth most common diagnosed cancer and the second leading cause of cancer-related deaths. This surge in liver cancer incidence is primarily driven by the two major subtypes, HCC and CCA, which together account for 95% of all cases. Detecting liver cancer early remains difficult due to its asymptomatic nature until advanced stages [1–3]. Challenges are exacerbated for CCA due to various environmental and socioeconomic factors that vary globally. Despite limited detection tools, risk factors like viral infections and metabolic disorders have been associated with increased CCA risks [4–8]. CCA, a rare cancer arising from bile duct epithelium, is classified into intrahepatic (iCCA), perihilar (pCCA), and distal (dCCA) based on location [6,7,9–11]. Despite its rarity, CCA constitutes 2% of all cancers, with rising occurrences in Western countries [2–4,6,7,10–14]. Inflammation is considered a significant contributor to CCA metastatic growth [15,16]. Incidence is influenced by genetics, risk factors, and regional disparities, with higher rates in the Eastern hemisphere and recent research explored the use of proteins from serum for non-invasive diagnosis and proposes targeting specific receptors for therapeutic strategies.[2–4,14,17].

Generally, cancer stands out as one of the most deadly illnesses in recent history, leading to numerous fatalities annually. As scientific progress continues, a range of medications and diagnostic approaches have surfaced to manage different types of cancer, contributing to the partial success in treating this ailment [1,18–20]. Identifying and documenting patients with cirrhosis and assessing liver masses are essential for effective CCA prevention, patient care and management [21]. Serum proteins from extracellular vesicles can aid non-invasive diagnosis [4,5]. Therefore, identification and development of biomarkers are important steps in the early screening, diagnosis, therapeutic evaluation, recurrence and prognosis prediction of tumors [22–24]. Omics is a powerful tool utilized to identify CCA-subtype specific biomarkers [19,20,25,26].

Cancer involves uncontrolled cell division caused by disrupted cellular transcription regulation [27]. Hence, mutated or improperly regulated TFs in different cancers serve as clear targets for therapeutic drugs as well as diagnostic and prognostic biomarkers [28,29]. Numerous new prognostic biomarkers or biomarker combinations derived from TFs have been identified through comprehensive omics analyses of transcription profiles across different cancer types [30–34], including HCC [31,35]. As numerous cases of liver cancer arise from chronic health conditions like cirrhosis triggered by hepatitis B or hepatitis C infections [18–20,22,23,36–38], alterations in gene expression induced by viruses have been documented at both the DNA and RNA levels [38–42]. This underscores the importance of TFs participating in the development of liver cancers. As a result, one objective of this research is to pinpoint the TFs associated with the potential diagnostic and prognostic biomarkers identified in CCA.

The public repositories housing liver cancer-related data have rapidly expanded, now encompassing a wide array of comprehensive information, including epidemiology, clinical details, diagnostic data, and Omics data[43–48]. The upward trend in liver cancer data repositories is anticipated to persist, continually providing new insights to enhance the existing diagnostic and prognostic biomarker collection for liver cancer, which is presently lacking. This study aims to identify diagnostic and prognostic biomarkers for CCA through the utilization of bioinformatics and data sourced from public repositories. By conducting a comparative analysis, we discovered potential gene biomarkers linked to the age and survival of individuals with CCA. Through pathway analysis, we unveiled functions and disrupted pathways correlated with these biomarkers, providing insights into the progression of the disease. Finally, elevated CCM protein levels in tumors night suggest their viability as biomarkers across multiple cancers [49–59]. Both main types of progesterone receptors (nPRs and mPRs) might serve as potential biomarkers for genetically influenced diverse cancer subtypes [60–68]. In this research, the utilization of publicly accessible data and our previous findings, we aim to conduct bioinformatics analysis on public datasets combined with our data to uncover potential prognostic biomarkers for CCA, thus enhancing our comprehension of molecular profiles and clinical importance in liver cancer, particularly CCA.

## Materials and Methods

### CCA patient data

Patient data for individuals with hepatic cancer was gathered from the openly accessible liver tissue data provided by The Cancer Genome Atlas (TCGA). A comprehensive set of 470 patient records was amassed, and these samples were thoroughly examined to establish their identities in relation to the subtype of hepatic cancer. Subsequently, our focus was on identifying 42 patient transcriptome profiles that had been diagnosed with a specific form of hepatic cancer known as CCA. Consequently, our analysis leaned towards the prognosis/diagnosis of CCA patients. This involved utilizing the number of days until either the patient’s demise due to the disease (20 cases) or the survival of the patient (22 cases) as the basis for assessment.

### Transcriptomic Analysis of CCA Subtype Using TCGA Data

To understand the duration of the disease, survival curves were employed, which represented the probability of survival over time. Our investigation into patient transcriptomes persisted as we meticulously gathered data on RNA sequences encoding proteins. This kind of data was vital in capturing alterations in functional units that were influenced by the progression of cancer. The survival outcome of each patient was categorized and used as a prognostic indicator. This allowed us to compare the gene expressions of over 19,000 genes, measured in transcripts per million (TPM), between those patients who survived and those who succumbed to the disease. To establish potential prognostic biomarkers, a Welch’s t-test was utilized to determine the significance of differences in means between the two groups. The results of this analysis enabled us to pinpoint potential prognostic biomarkers linked to patient outcomes and the expression of tissue protein coding RNA sequences.

### Normal liver tissue controls for comparative analysis of prognostic biomarkers

Furthermore, apart from identifying the prognostic biomarkers, we procured control samples of liver tissue from the GTEx portal. This yielded a total of 272 control liver tissue samples. From these, we extracted the protein-coding RNA sequences that had been identified as prognostic indicators. To compare, we comparatively selected the top 25 statistically significant genes from the group of patients with CCA against the expression profiles of these control samples from healthy individuals.

### Pathway analysis for candidate prognostic biomarkers with Gene Expression data

To the total number of genes that displayed statistically significant differences between the survivor and deceased CCA patient cohorts. We performed gene set enrichment analyses (GSEA) on genes displaying the highest level of statistical significance. To gain a comprehensive understanding of the pathways impacted and contributing to distinct prognosis, we conducted the GSEA analysis across three distinct libraries (GO, KEGG, and DOSE). Continuing our inquiry, we implemented a filtering mechanism specifically targeting genes exhibiting unique expressions linked to HCC. This approach aimed to exclude genes that were shared between both cancer types. As a result, a total of 24 such genes, which held significance in influencing patient prognostic outcomes, were eliminated. Following that, a secondary GSEA analysis on genes displaying the highest level of statistical significance was carried out, specifically aimed at identifying pathways that were uniquely affected by CCA.

### Statistical analysis

Through the application of the t-test, we investigated not only the significantly differential expressions between the healthy control and CCA cohorts but also delved into the significantly differential expressions within the survival and deceased CCA patient groups, subsequently comparing them to the control group. To analyze the differentia expression patterns of protein coding RNAs, we initially employed a Welch’s t-test to identify TPM values of mRNAs. This outcome facilitated the identification of differences in prognostic status, leading us to establish a filter for mRNAs displaying differential expression. Subsequently, we exclusively compared the statistically significant differentially expressed mRNAs identified in the initial step, contrasting CCA survivors with the normal controls. This analysis was conducted to ascertain the significance of the differential expression in relation to the control population. Our objective was to pinpoint exclusively distinct TPM values to establish diagnostic potential. In conclusion, we employed another Welch’s t-test on mRNAs showing significant differences between CCA survivors and controls. Next, we aimed to determine if these protein coding mRNAs exhibited significant differential expression in the deceased CCA cohort. This approach enabled the identification of unique mRNAs that showcased differential expression not only in survival and deceased CCA cases, but also in both cohorts when compared to controls. This established both prognostic and diagnostic capabilities (Figure 2A). Alongside the establishment of diagnostic and prognostic biomarkers, another aim of this study encompassed gauging the differential expression pattern between CCA patients and the healthy controls. To achieve this objective, we utilized heatmaps, which visually portrayed the number of differential expressed genes (DEGs) (Fig. 3). Finally, expanding beyond the list of 25 statistically significant DEGs, our investigation extended to try to incorporate members of the CmPn signaling network, due to the potential relevance of liver steroid hormones in our study context [64].

### Transcription Factor prediction and Experimental Setup

The challenge associated with predicting transcription factors (TFs) arises from the intricate and costly process of experimental verification. In this study, we introduce the ESM-TFpredict framework aimed at automating the identification of potential TFs. ESM-TFpredict exploits the advanced ESM-2 pre-trained protein language model to represent protein sequences [69,70]. Subsequently, it employs 1-D Convolutional Neural Networks (CNNs) to predict transcription factors [71]. Based on the architecture of ESM-TFpredict (Suppl. Figure 1), the process commences by assigning a semantic representation to every amino acid present in the protein sequence, employing the pre-trained ESM-2 model. To optimize training efficiency while adhering to limited resource availability, the sequence is divided into predetermined windows. The amino acid representations within each window are averaged to create a concise representation. The prediction model then utilizes two layers of 1-D CNNs (with 128 and 256 filters respectively), each followed by a max pooling and dropout layer (with a dropout rate of 0.5). Subsequent to these layers, a dense layer and another dropout layer are applied to further refine the extracted features and counteract overfitting. Ultimately, the final prediction is generated through sigmoid activation.

#### Dataset Preprocessing

Our protein dataset was mainly obtained from the UniProt (www.uniprot.org). To assemble a collection of TF sequences with high confidence, we sifted through the Swiss-Prot dataset, focusing on annotations containing the keywords “transcription regulation” and “DNA binding” [72]. This curation yielded a total of 5,418 TF sequences, all of which carried a perfect 5/5 annotation score. For sequences not associated with transcription factors (referred to as NTF sequences), we intentionally excluded instances containing the aforementioned keywords, which resulted in a set of 47,120 sequences. In order to establish dataset equilibrium, we randomly selected 5,564 NTF samples. It is noteworthy that more than 98% of the sequences have a length of under 3,000 amino acids (Suppl. Table 1). To ensure uniform analysis, we will cap the sequence length at 3,000 in our experiments, truncating longer sequences accordingly.

#### Experimental execution and evaluation

The experiments were performed utilizing a Linux server equipped with an Intel(R) Core(TM) i9-10900X CPU operating at 3.70GHz. This system was further enhanced by a GeForce RTX 3080 GPU and boasted a RAM capacity of 122GB. During the training of the ESM-TFpredict model over 40 epochs, a batch size of 128 was employed alongside a learning rate of 10^-4. The condensed representation was standardized at a fixed size of 300. By employing 5-fold cross-validation, we acquired outcomes that encompassed an average F1 score of 0.9531, specificity measuring 0.9537, sensitivity at 0.9525, and a balanced accuracy reaching 0.9531. These scores emphasize the dependability and precision of our machine learning (ML)/deep learning (DL)-based TF prediction model.

## Results

### Age Distribution, Survival Trends, and Prognostic Possibilities within our CCA Patient

At the outset of our investigation utilizing the TCGA-derived dataset, we assembled a group comprising 42 patient profiles afflicted with CCAs. Initially, we sought to delineate the age distribution within this cohort and decipher the patterns associated with survival rates (Figure 1A). Then, we examined CCA patient life span, visualized by the KM survival curves (Figure 1B). Our data unveiled a discernible trend: with the passage of time, the overall prognosis linked to CCA diagnosis experienced a swift deterioration (Fig. 1). The overall prognosis of CCA diagnosis rapidly worsened over time. Notably, none of the patients survived beyond the 2000-day point with an ongoing diagnosis, aligning with previous research highlighting the inherently unfavorable CCA prognosis [73,74].

**Figure 1.**
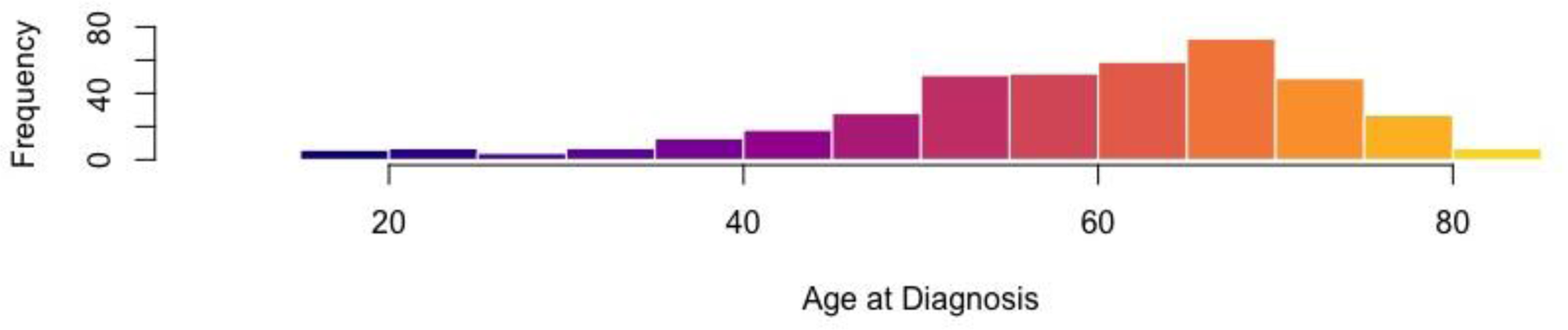
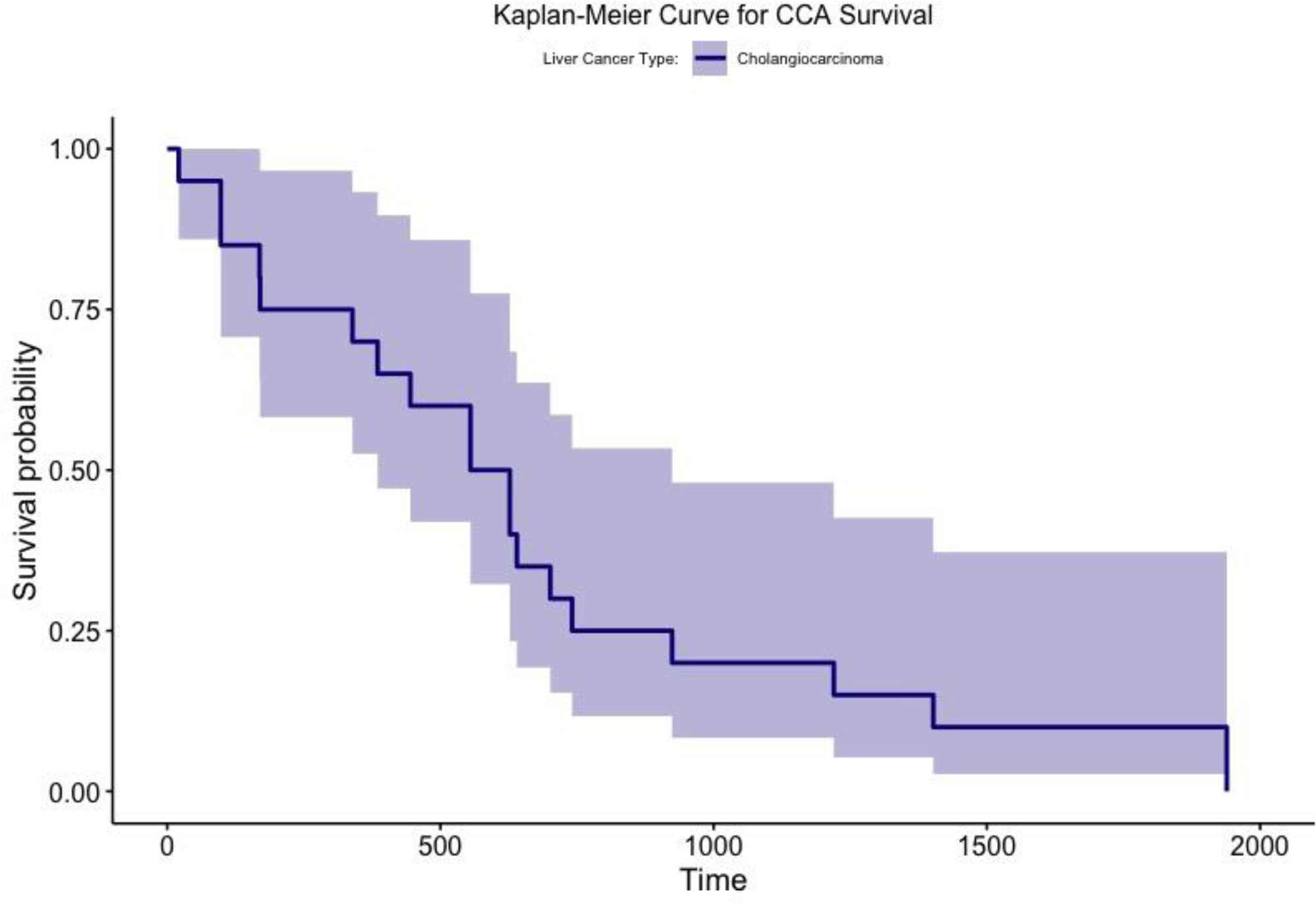
Demographic Analysis and Survival Patterns of Cholangiocarcinoma (CCA) Patients Based on TCGA Data. Liver cancer represents an intricate ailment with limited comprehension concerning its timely identification. Among its various subtypes, the second most prevalent form is Cholangiocarcinomas (CCAs). Scrutinizing the data sourced from TCGA, we collected 42 patient profiles afflicted by CCAs. Our intention was to examine the demographic distribution of ages within this group and ascertain the patterns of survivability. A. The patient cohort encompassed those diagnosed with CCA subtype. The age spectrum spanned from the mid-teenage years to eighty-five. Notably, the highest vulnerability to diagnosis was observed between the fifth and eighth decades of life. B. Kaplan-Meier Survival Analysis of CCA Patients revealed that none of those with CCA who succumbed to the illness survived beyond 2000 days.

### Identification of Prognostic and Diagnostic Biomarkers from Comprehensive Analysis of Protein-Coding RNA Sequences in CCA Patients cohort

Our subsequent investigation encompassed a comprehensive examination of over 19,000 protein-coding mRNAs, revealing 388 potential candidates with significant prognostic relevance. These candidates displayed varying transcript per million (TPM) expressions in relation to survival outcomes. Upon further analysis of the 388 prognostic candidates we furthered refined the results by checking for statistical difference not only between the CCA survivors and the normal controls, but also between the deceased CCAs and normal controls (Fig. 2A).

**Figure 2.**
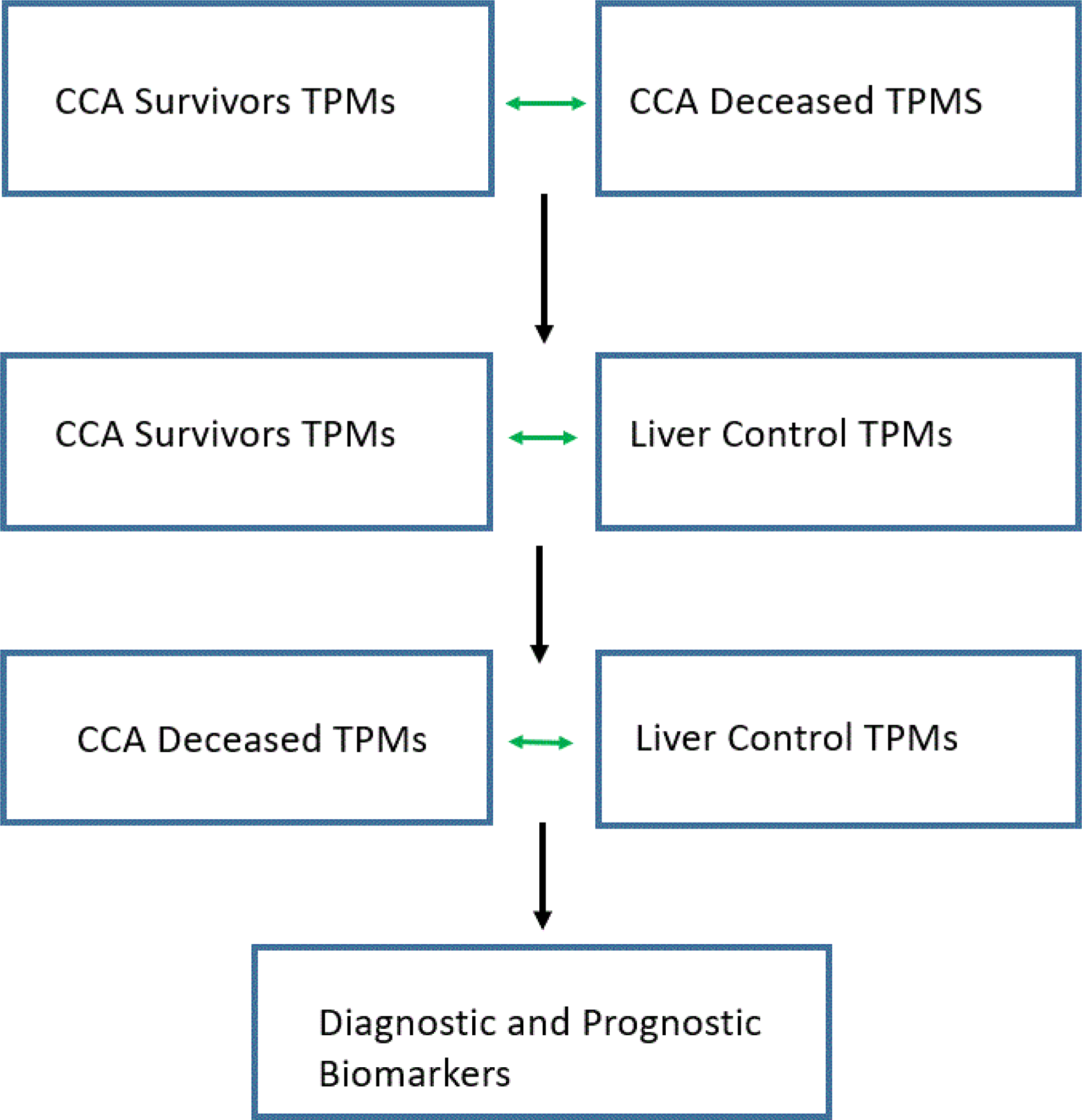
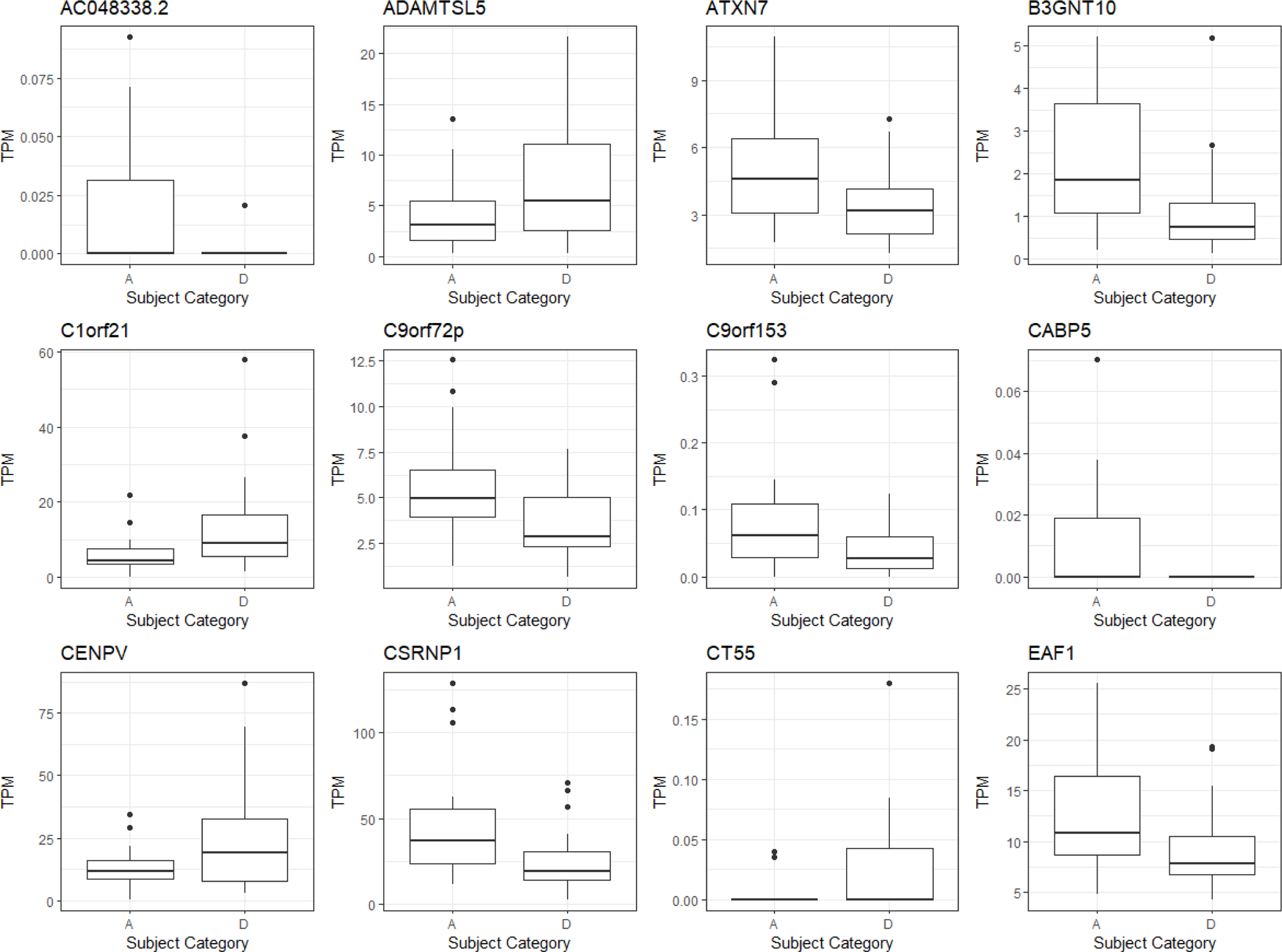
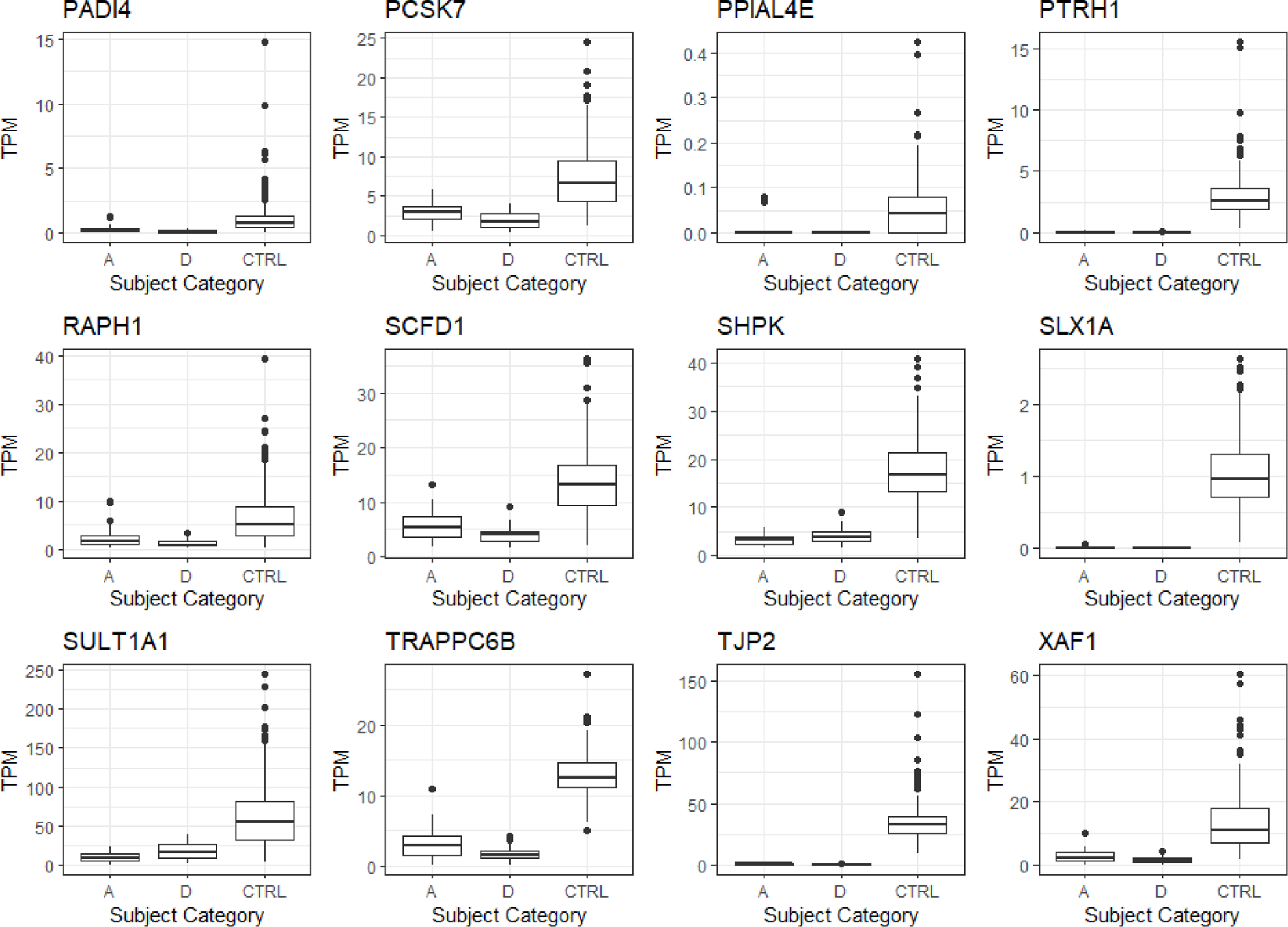
Prognostic Biomarker Potential of Protein-Coding Genes in CCA. Boxplots were utilized to illustrate the dispersion of data concerning protein-coding genes potentially functioning as prognostic biomarkers. These genes’ expressions, quantified by Transcripts per Million (TPM), were categorized by the samples’ vital status. Sourcing the data from the TCGA database, the TPM values for Control (CTRL), survivors (A), and deceased (D) samples were collated. It’s noteworthy that all the genes displayed in the plots are plausible candidates for prognostic biomarkers. **A**. This diagram portrays the flowchart of the analysis and experimental design. Each box represents a distinct analytical step corresponding to the types of datasets generated within each cohort. **B.** Box plots were employed to visualize the top results of protein-coding genes in CCA, chosen based on the most significant statistical distinctions among the alive, and deceased samples. These genes exhibit potential prognostic biomarkers. (Note: The second part of these plots is illustrated in Supplemental Figure 2B) **C.** Box plots were employed to visualize the top results of protein-coding genes in CCA, chosen based on the most significant statistical distinctions among the alive, deceased, and control samples. These genes exhibit potential as both prognostic and diagnostic biomarkers. (Note: The second part of these plots is illustrated in Figure 2C)

For a more comprehensive investigation, our focus was directed towards the 388 differentially expressed candidates. These candidates not only displayed substantial differential expressions between normal controls and CCA patients who survived, but also exhibited significant differences in their expression pattern between surviving and deceased CCA patients (Figs 2B-C, Suppl. Figs 2B-C). These differential expression patterns in CCA patients were further visualized using heatmaps (Fig. 3). Based on these findings, we concluded that the potential exists for identifying both prognostic-exclusive biomarkers and biomarkers with combined prognostic and diagnostic capabilities (Figs. 2, 3, Suppl. Fig. 2). In fact, this two-phase analysis highlighted a total of 187 promising candidates for prognosis and diagnosis (Table 1).

**Figure 3.**
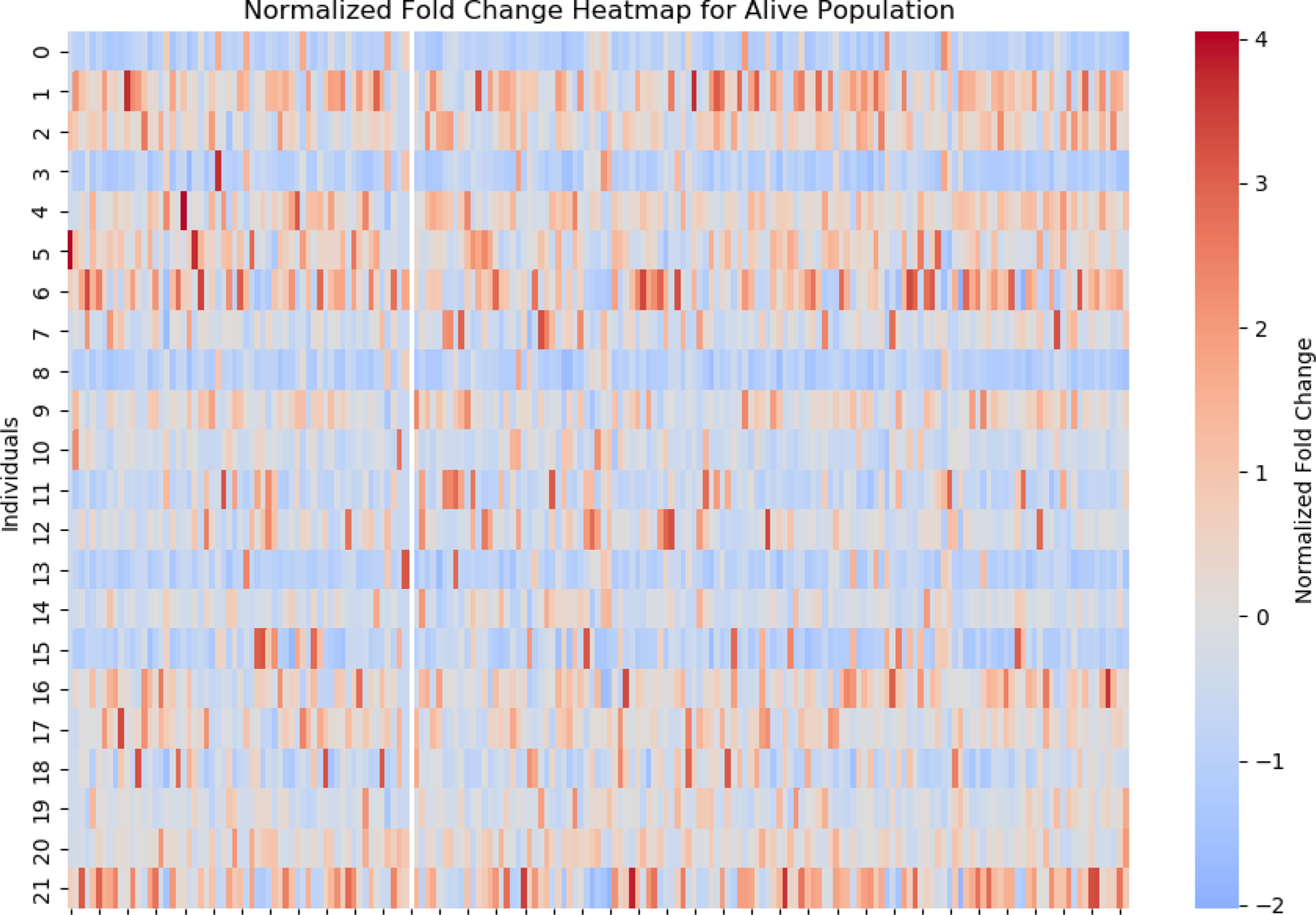
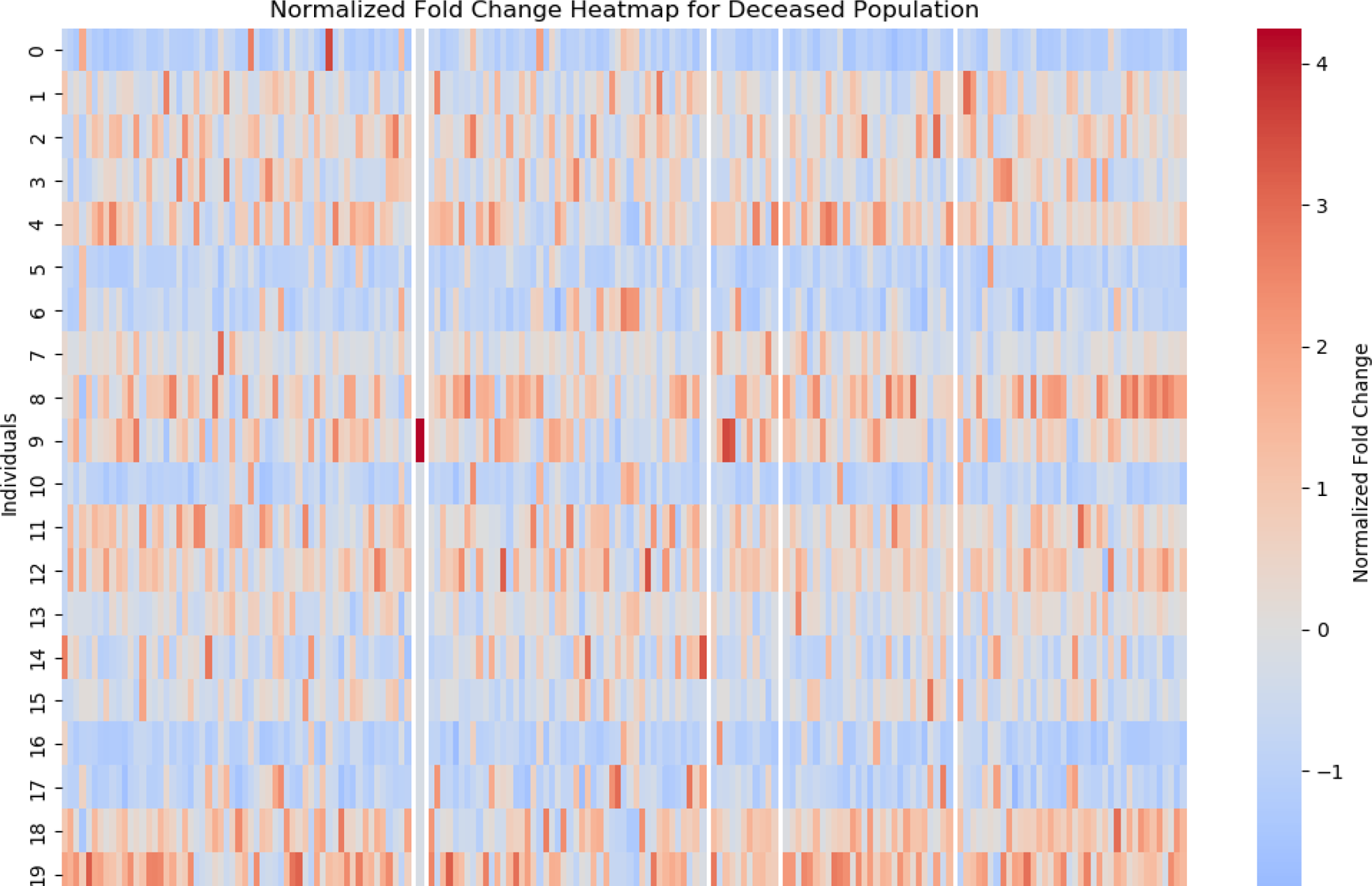
Identification of CCA-specific prognostic biomarkers from altered signal pathways through comparative enrichment Analysis between CCA patients and healthy controls. Apart from previously identifying distinct prognostic and diagnostic biomarker candidates, we conducted a comparison against the normalized control values to determine the fold change values. These values are visually represented through the use of heatmaps. **A**. The analysis illustrates the normalized fold change values for protein-coding RNA sequences within the survivor population. We computed the change in TPMs to present the fold change analysis. Notably, a significant portion of the expressions in the survivor group exhibited upregulation across the 187 coding RNA sequences. Patient identifiers are depicted on the y-axis, while the 187 coding RNA species are represented on the x-axis. **B.** Similarly, this analysis exhibits the normalized fold change values pertaining to the deceased patients. In contrast to the survivors, fewer pathways display upregulated genes. The labeling scheme on the x and y-axes corresponds to the 187 coding RNA sequences and the patients, respectively.

**Table 1.**
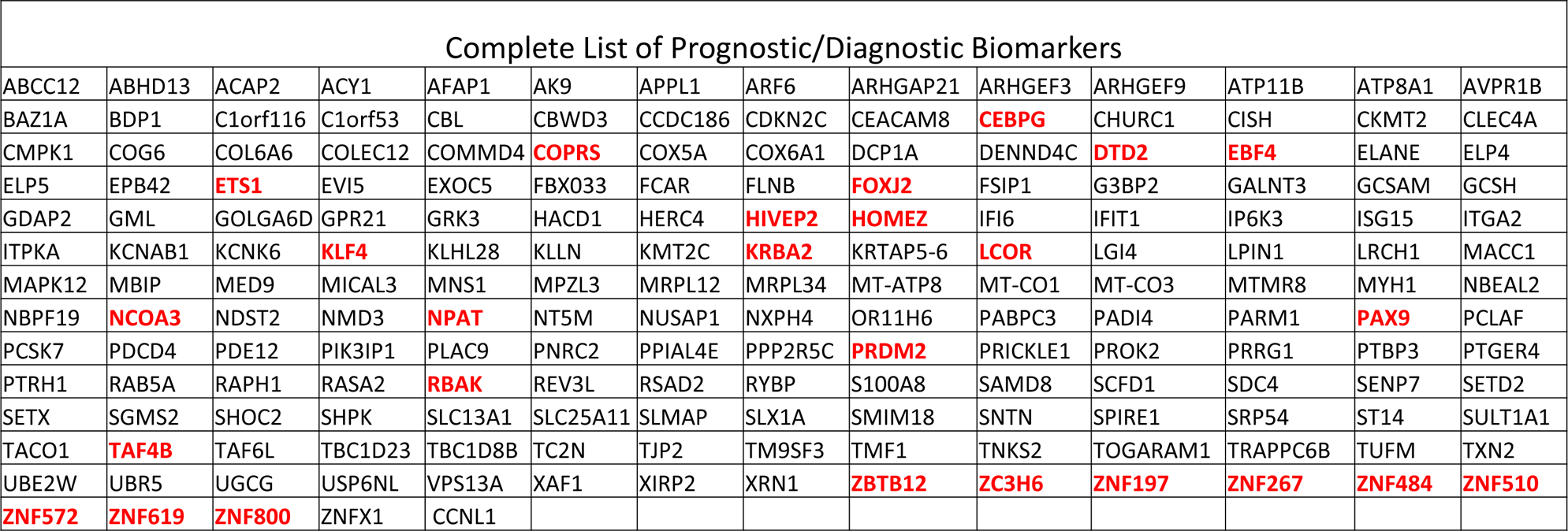
The complete list of the prognostic and diagnostic biomarkers identified in this study with all statistical significance. Comprehensive list of Prognostic and Diagnostic Biomarkers Identified through an SVM model, inclusive of All Statistical Significance. The gene symbols shown in red color above correspond to RNA sequences that machine learning identified as potentially having TFs activity through an SVM model.

Concurrently, we investigated the viability of members within the CmPn signaling network as potential biomarkers. Our findings indicated that, although none of the CmPn members fulfilled the role of prognostic biomarkers for this specific subtype of hepatic cancer, however, we found that out of the 14 CmPn network members, 9 had the potential to be employed as diagnostic tools (Suppl. Fig. 2A).

### Discovery of Prognostic Biomarker through Pathway Enrichment in CCA

Next, we embarked on a series of comprehensive pathway enrichment analyses aimed at understanding the functional implications of genes with significantly differential expression as prognostic biomarkers unique to the CCA. This study involved employing Gene Set Enrichment Analyses (GSEAs) across three distinct libraries (GO, KEGG, and DOSE), specifically tailored to the genes recognized as unique prognostic biomarkers in CCA (Suppl. Figs. 3A-C). By conducting these analyses, our aim was to reveal the biological pathways and molecular functions linked to these genes through a combined comparative GSEA approach. This shed light on their potential roles within the context of CCA prognosis (Fig. 4A).

**Figure 4.**
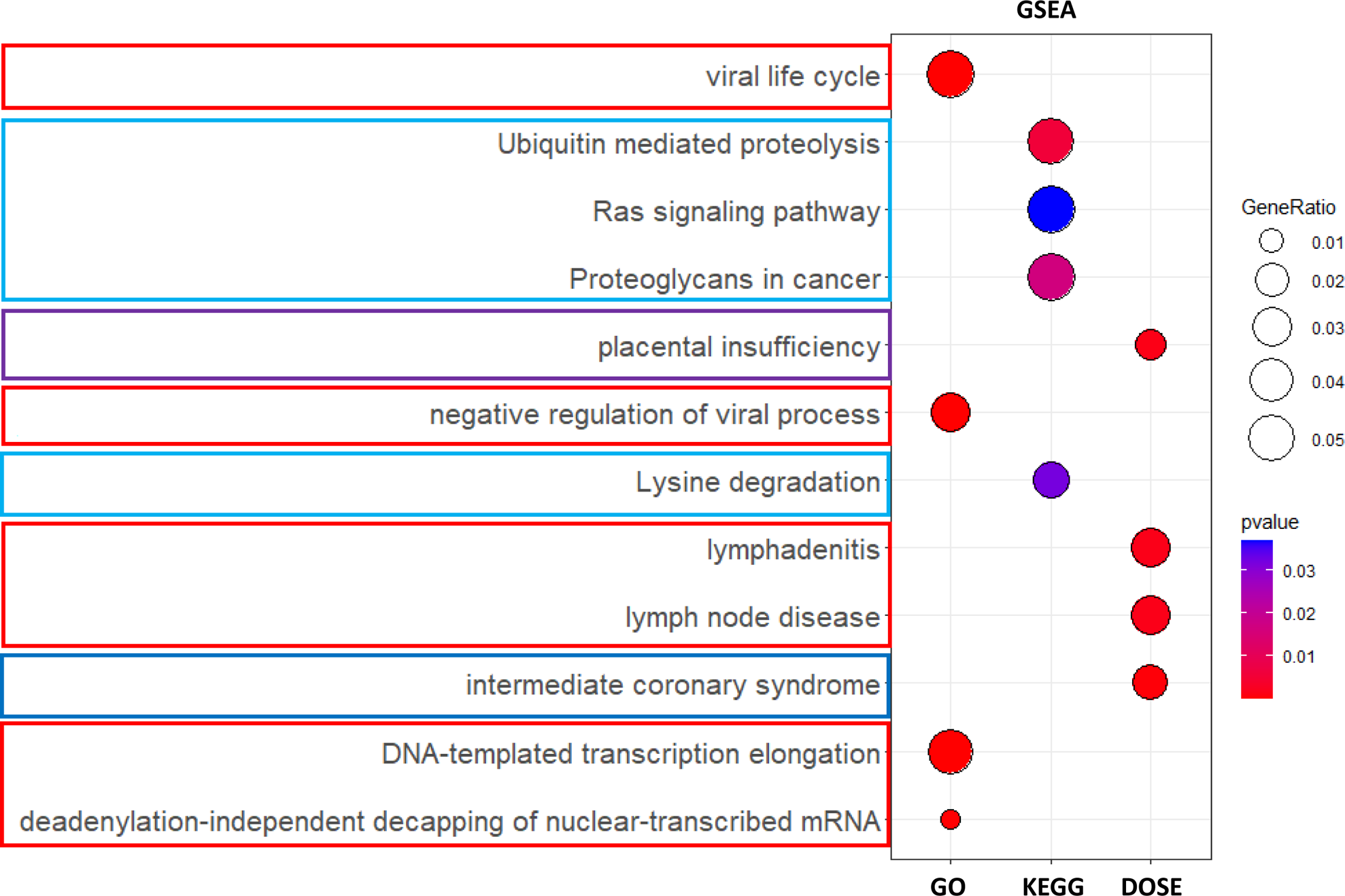
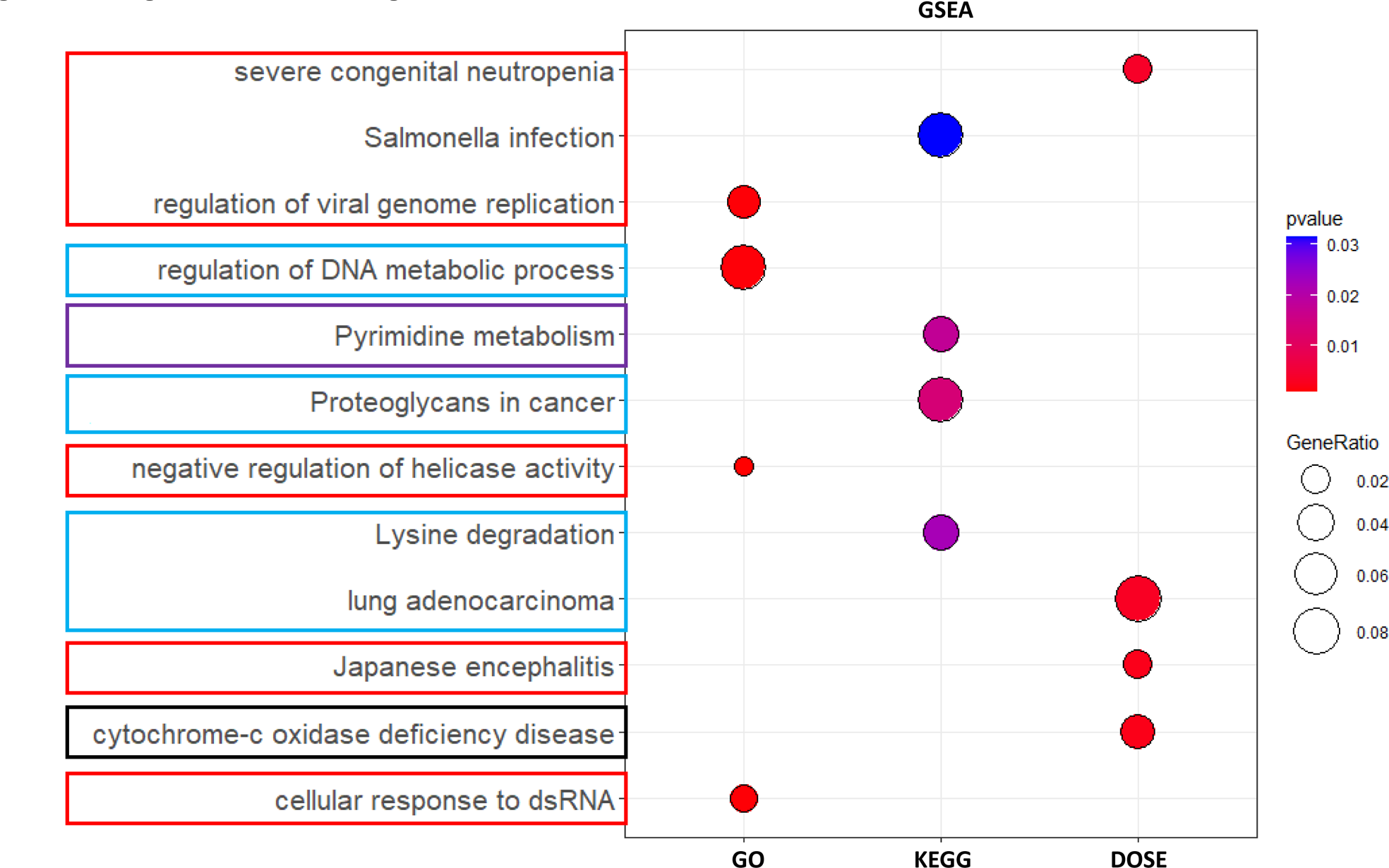
Uncovering unique Cholangiocarcinoma (CCA) prognostic biomarkers through Gene Set Enrichment Analyses (GSEA). By utilizing three separate enrichment libraries, we highlight the pathways influenced by CCA tumorigenesis, with the intention of identifying the primary diseases associated with these genes. **A.** This plot visualizes the signal pathways linked to 388 identified RNA species, which have been identified as potential prognostic biomarkers through a comparative expression analysis conducted between surviving and deceased CCA patients. **B.** Similarly, this plot depicts the signal pathways linked to 187 identified RNA species, which have been recognized as potential diagnostic and prognostic biomarkers. This recognition is based on a comparative expression analysis conducted among survival CCA patients and controls, as well as deceased CCA patients and controls. Moving across from the left, we can see enriched analyses involving gene ontology (GO), Kyoto Encyclopedia of Genes and Genomes (KEGG), and Disease Ontology Semantic and Enrichment (DOSE). These analyses highlight pathways impacted by viral infections and inflammatory responses (indicated by red outlines), tumorigenesis (light blue outline), hormonal responses (purple outlines), and metabolic pathways (outlined in black). The pathways that overlap between these two analyses offer promise for identifying diagnostic and prognostic biomarkers, which can be subjected to further validation analysis in the future.

### Identification of Dual diagnostic and prognostic biomarkers and Associated Pathways in CCA via Comprehensive GSEA

We then proceeded to perform an enrichment analysis aimed at identifying the signal pathways that were most profoundly disrupted by the CCA. This analysis entailed a meticulous comparison between pathways deemed normal and those exhibiting significant statistical differences within both surviving and deceased CCA patient groups. Additionally, we thoroughly examined the statistical significance of gene expression patterns across normal controls and distinct groups of surviving and deceased CCA patients. This investigation encompassed pathway enrichments spanning three distinct libraries (GO, KEGG, and DOSE) to define CCA-specific diagnostic and prognostic biomarkers (Suppl. Figs. 4A-C). The ultimate comprehensive results enabled us to identify the particular signaling pathways substantially impacted by the tumorigenesis of CCA. This process also facilitated the recognition of intricate molecular alterations linked to the tumorigenesis and progression of CCA, achieved through a combined GSEA analysis. (Fig. 4).

### Some diagnostic/prognostic biomarkers act as TFs to regulate Associated Pathways in CCA

By employing our newly developed ML/DL-based ESM-TFpredict model (Suppl. Fig. 1, Suppl. Table 1), we have successfully discerned 26 transcription factors (TFs) from the 187 diagnostic and prognostic biomarkers specific to CCA that we had previously discovered (Table 1, red-colored). This discovery is consistent with the transcription profiling data reported in various cancer types [30–34], including HCC [31,35]. Recognizing the significant role of TFs in the progression of liver cancers [38–42], we conducted enrichment analyses to unveil the signaling pathways in which the identified TFs are implicated in the context of CCA. By utilizing three enrichment libraries (GO, KEGG, DOSE), our analyses were directed towards the identification of CCA-specific TFs with potential as prognostic biomarkers (Suppl. Figs. 5A-C). The effort to delineate altered signaling pathways in CCA associated with these TFs has been successfully realized through a combined GSEA assessment (Fig. 5). Intriguingly, three of the four main signal pathways, encompassing inflammatory responses (red outlines), tumorigenesis (light blue outline), and hormonal responses (purple outlines), exhibit an overlap with signal pathways previously defined within diagnostic and prognostic biomarkers (Figs. 3 and 4). This overlap strongly implies that these pathways are under the regulation of the identified TF biomarkers.

**Figure 5.**
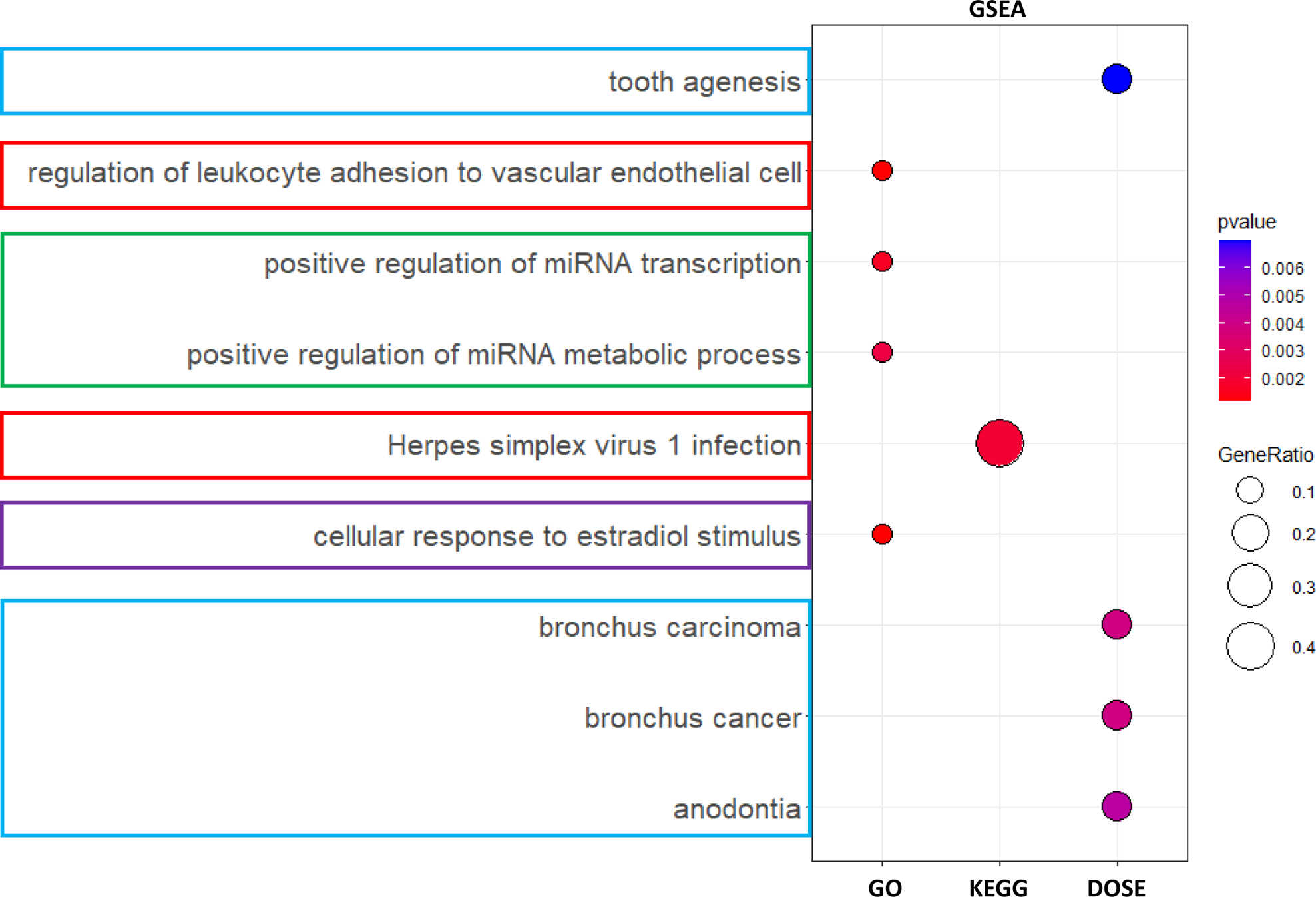
Uncovering Altered signal pathways in CCA through different Enrichment Analysis. Enrichment analysis were conducted to ascertain the communication pathways most significantly disrupted by CCA. This analysis involved comparing normal pathways to those that exhibited statistical differences within both the surviving and deceased populations, as well as between the groups of survivors and deceased individuals with statistically distinct gene expression patterns. This analysis shows the pathways that are enriched through three different libraries along the x-axis. The predicted transcription factors were found to affect the above pathways. Enrichment analyses were performed by employing three distinct enrichment libraries, aiming to pinpoint the CCA-specific TFs as prognostic biomarkers. Moving from left to right, we observe enriched analyses involving gene ontology (GO), Kyoto Encyclopedia of Genes and Genomes (KEGG), and Disease Ontology Semantic and Enrichment (DOSE). These analyses help identify the most significantly disrupted signal pathways in CCA, associated with these TFs. These analyses highlight pathways impacted by viral infections and inflammatory responses (indicated by red outlines), tumorigenesis (light blue outline), and hormonal responses (purple outlines). The color-emphasized pathways, which coincide with earlier GSEA results indicating these TFs as potential diagnostic and prognostic biomarkers, are believed to operate via transcriptional control mechanisms.

## Discussion

### Distinct clinical/diagnostic characteristics between HCC and CCA

There are two major groups of liver cancer based on common clinical/diagnostic criteria, either Hepatocellular carcinoma (HCC) or intrahepatic bile duct cancers (Cholangiocellular carcinoma, CCA) [18–20,22,23,36]. Recently, we conducted an initial comprehensive clinical analysis of these two liver cancers, with main focusing primarily on HCC [64]. This involved harnessing data from publicly accessible databases. Additionally, we extended our clinical investigation specifically to the members of the CmPn network (CCM1-3; PAQR5-9; and PGRMC1-2), leveraging two databases containing patient samples with differential expression data for these key CmPn components. Sociological, pathological, and follow-up clinical data [25] for HCC (and some CCA) patients, encompassing 438 HCC tumor samples, were extracted from the TCGA-LIHC and TCGA-PANCAN databases. These data were utilized for expression profiling, allowing us to explore differentially expressed genes across factors such as race, gender, family cancer history, vascular invasion, stemness scores across histological types, new tumor events (type and site), vital status, and residual tumor status [25].

### Comparative analysis reveals divergent trends in for biomarkers between HCC and CCA

HCC stands as the most prevalent form of primary liver cancers, frequently manifesting in individuals grappling with persistent liver conditions such as cirrhosis stemming from hepatitis B or hepatitis C infections [18–20,22,23,36]. Our prior research outcomes unveiled strong clinical connections between all components of the CmPn network in HCC, while only some members of CmPn network members with CCA [64]. Furthermore, comparative Omics unveiled a predominant down-regulation of CmPn genes in CCA, setting it apart from the up-regulation of CmPn genes in HCC [64]. This underscores the fundamental distinction between the two major liver cancer types [64].

This study aimed to identify biomarkers through bioinformatics analysis of publicly available datasets. The research focuses on CCA patients’ transcriptomes, aiming to identify potential gene biomarkers associated with CCA patient age and survival. Pathway analysis helps elucidate functions and disrupted pathways correlated with these biomarkers. The study also explores the prediction of transcription factors (TFs) using a machine learning model called ESM-TFpredict. This model automates the identification of potential TFs, leveraging protein sequence data. The dataset preprocessing involves curating TF sequences and non-TF sequences for analysis. The model’s experimental execution and evaluation demonstrate its reliability and precision in TF prediction. Overall, the study combines bioinformatics analysis, transcriptomic investigation, and machine learning to uncover potential diagnostic and prognostic biomarkers for CCA. The utilization of publicly available data enhances our understanding of liver cancer’s molecular profiles and clinical significance, particularly in the context of CCA.

