## Supplementary material for "Identification of Cholangiocarcinoma (CCA) Subtype-Specific Biomarkers": Suppl Materials

Running title: CCA Specific Biomarkers

**Supplemental Materials**

*All correspondence:*
Jun Zhang, Sc.D., Ph.D.

Department of Molecular & Translational Medicine (MTM)
Texas Tech University Health Science Center 5001 El Paso Drive, El Paso, TX 79905

**Supplemental Legends**

Supplemental Figure **1. The architecture of ESM-Tfpredict algorism.** The initiation of the ESM-Tfpredict procedure involves utilizing the pre-trained ESM-2 language model to allocate a semantic representation to each amino acid existing within the protein sequence. Subsequently, in order to enhance training efficiency while considering the constraints of available resources, the approach involves compacting representations through a process involving dual layers of 1-D CNNs (with 128 and 256 filters, respectively). These are sequentially accompanied by max pooling and dropout layers (with a dropout rate of 0.5) to generate final outcomes.

Supplemental Figure **2**. **Boxplot illustrate the prospective utility of proteins/genes within the CmPn pathway, and our identified prognostic biomarkers and prognostic/diagnostic biomarkers.** Boxplots were utilized to visualize the data distribution of protein-coding genes that could potentially act as diagnostic biomarkers. These genes' expression levels, measured in Transcripts per Million (TPM), were organized based on the vital status of the samples. The TPM data, obtained from the TCGA database, was collected for Control (CTRL), survivor (A) and deceased (D) CCA patient samples. **A.** By employing box plots, we illustrated elements of the CmPn signaling pathway with the potential to serve as diagnostic biomarkers. **B**. We continued to employ boxplots to finish the top results found in Figure 2B of the main figure legends. These results are the remaining top results of the prognostic biomarkers. **C.** The last of the top 25 diagnostic and prognostic biomarkers. This is a continuation of Figure 2C of the main figures.

Supplemental Figure **3**. **Exploring Enriched Prognostic Biomarkers through Gene Set Enrichment Analyses (GSEAs) in CCA.** Multiple pathway enrichment analyses were performed on genes with significantly altered expression patterns that were distinctly identified as exclusive prognostic biomarkers for CCA. These Gene Set Enrichment Analyses were exclusively carried out for genes recognized as unique prognostic biomarkers in the context of CCA. **A.** The application of Gene Ontology (GO) analysis was utilized to unveil enrichment patterns specific to prognostic biomarkers relevant only to CCA. The objective was to discern the biological processes, cellular compartments, and molecular functions influenced by these genes. **B.** Through enriched Kyoto Encyclopedia of Genes and Genomes (KEGG) analysis, we investigated genes identified as distinct prognostic biomarkers for CCA, with the intention of elucidating the key pathways affected by these genes. **C.** By utilizing enriched Disease Ontology (DOSE) analysis, we explored prognostic biomarkers exclusive to CCA. The goal was to differentiate the diseases primarily linked to these genes.

Supplemental Figure **4. Uncovering Diagnostic and Prognostic Biomarkers linked to Modified Signaling Pathways in CCA via Enrichment Analysis.** Pathway Enrichment analysis was executed to determine the signal pathways that were most significantly perturbed by CCA. This analysis entailed comparing regular pathways with those that displayed statistically significant differences within the groups of survival and deceased CCA patients, as well as between the groups of survival and deceased CCA patients with statistically differential gene expression patterns. **A.** GO enrichment analysis was employed to investigate the impact of CCA-specific prognostic biomarkers on biological processes, cellular locations, and molecular functions. **B.** Through KEGG enrichment analysis, we examined genes recognized as CCA-specific prognostic biomarkers, in order to identify the pathways that were most significantly impacted by these genes. **C.** Utilizing DOSE enriched analysis, we investigated prognostic biomarkers specific to CCA in order to establish the diseases that are most strongly linked to these genes.

Supplemental Figure **5. Unveiling TF-Regulated Pathway Alterations in CCA via Enrichment Analysis.** Enrichment analysis was conducted to identify the signal pathways regulated by the identified TFs, which exhibited the most notable disruptions due to CCA tumorigenesis. This analysis entailed comparing regular pathways with those that displayed statistically significant differences within the groups of survival and deceased CCA patients, as well as between the groups of survival and deceased CCA patients with statistically differential gene expression patterns. **A.** GO enrichment analysis was employed to investigate the impact of CCA-specific prognostic biomarkers on biological processes, cellular locations, and molecular functions. **B.** Through KEGG enrichment analysis, we examined genes recognized as CCA-specific prognostic biomarkers, in order to identify the pathways that were most significantly impacted by these genes. **C.** Utilizing DOSE enriched analysis, we investigated prognostic biomarkers specific to CCA in order to establish the diseases that are most strongly linked to these genes.

Supplemental Table 1: **The architecture of the transcription factor training model. The training model was approximately equal parts known transcription factors to non-**

**transcription factor FASTA sequences.** The model was trained on several different length sequences to have the widest range of learned transcription factors. This was applied to the machine learning model prior to the prediction of the final set of protein coding sequences.

Supplemental Figure 1. The architecture of ESM-Tfpredict algorism.

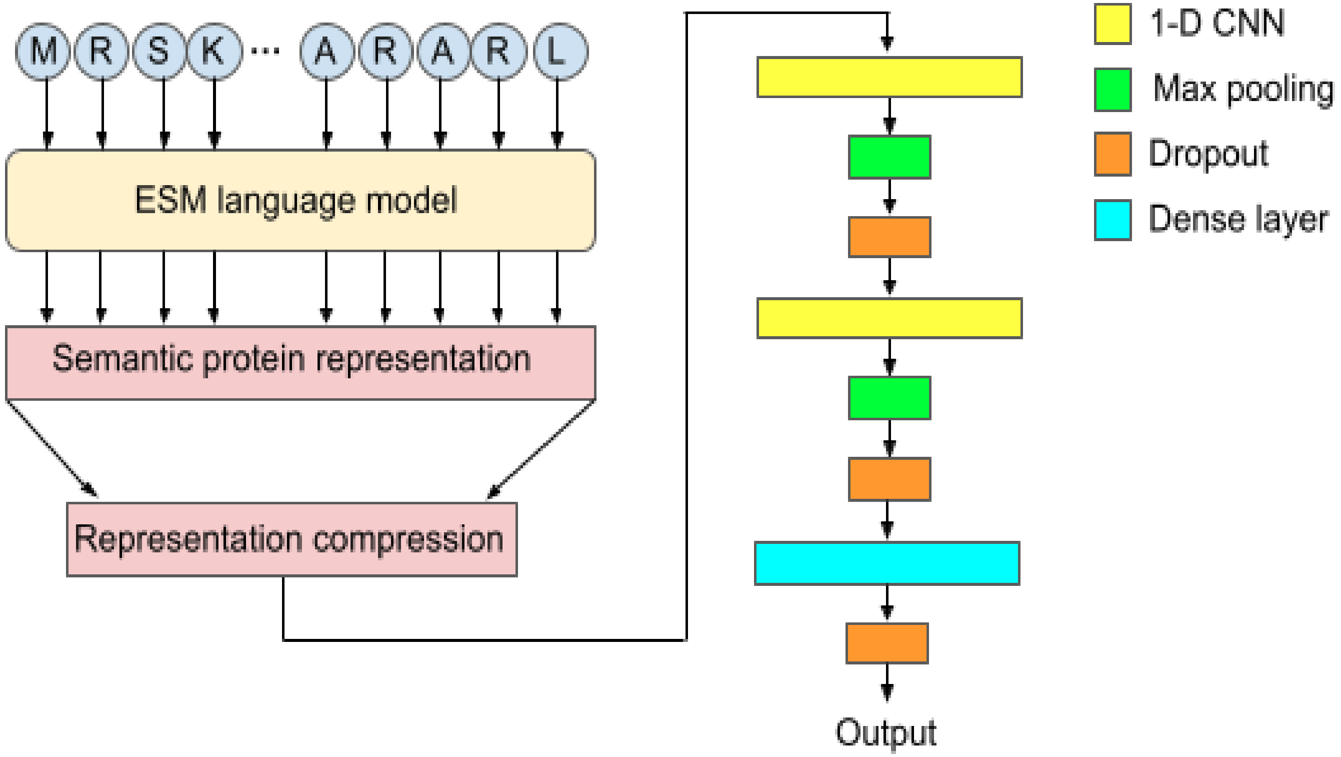

Supplemental Table1. Dataset distribution

| Sequence Length | [1,1000) | [1000,2000) | [2000,3000) | [3000, ) | Total |
|---|---|---|---|---|---|
| TF | 4786 | 521 | 85 | 26 | 5418 |
| NTF | 4628 | 713 | 134 | 89 | 5564 |

Supplemental Figure 2A: CmPn Members expression in CCA

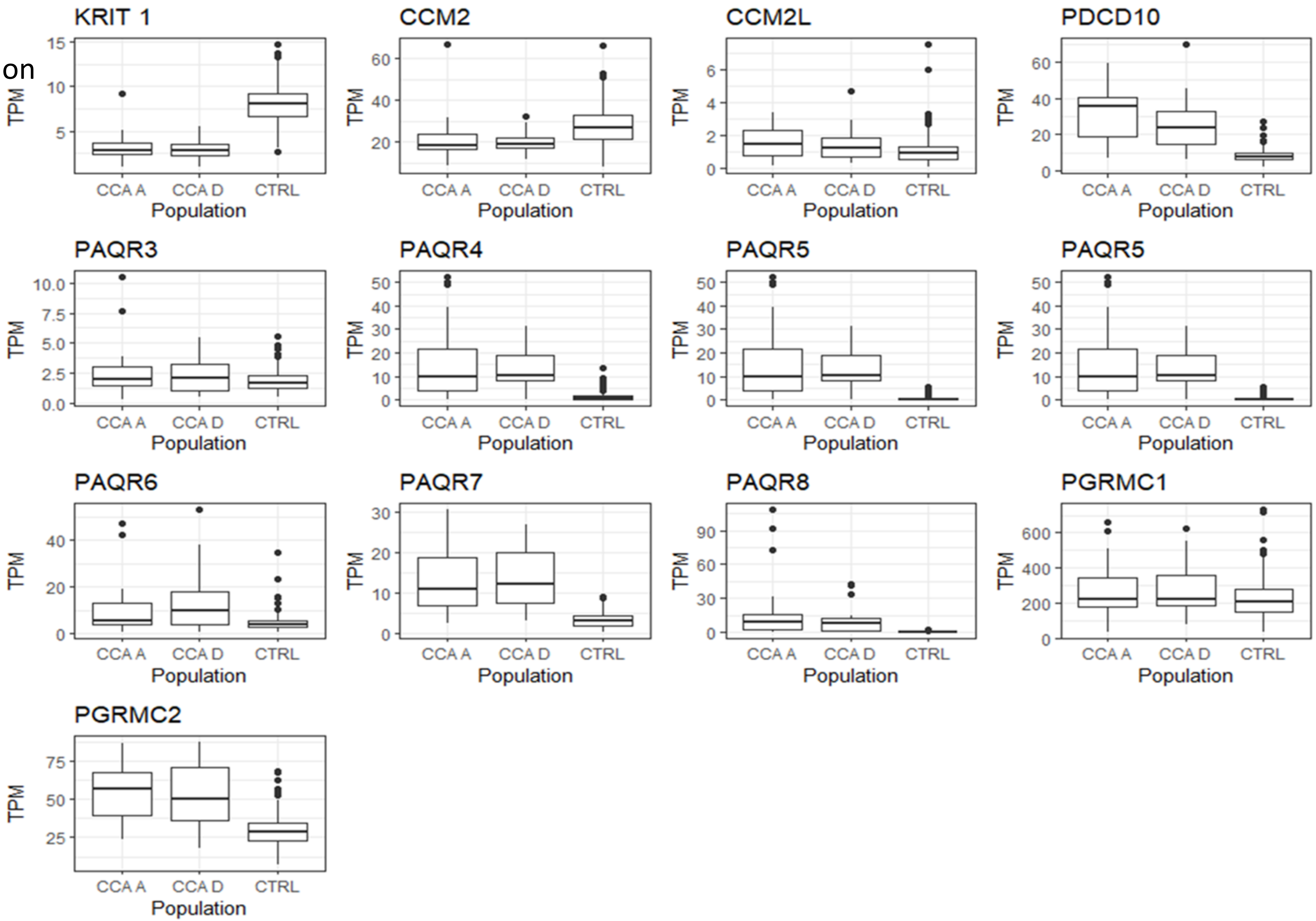

Supplement Figure 2B: Top Results of most significant prognostic biomarkers part 2

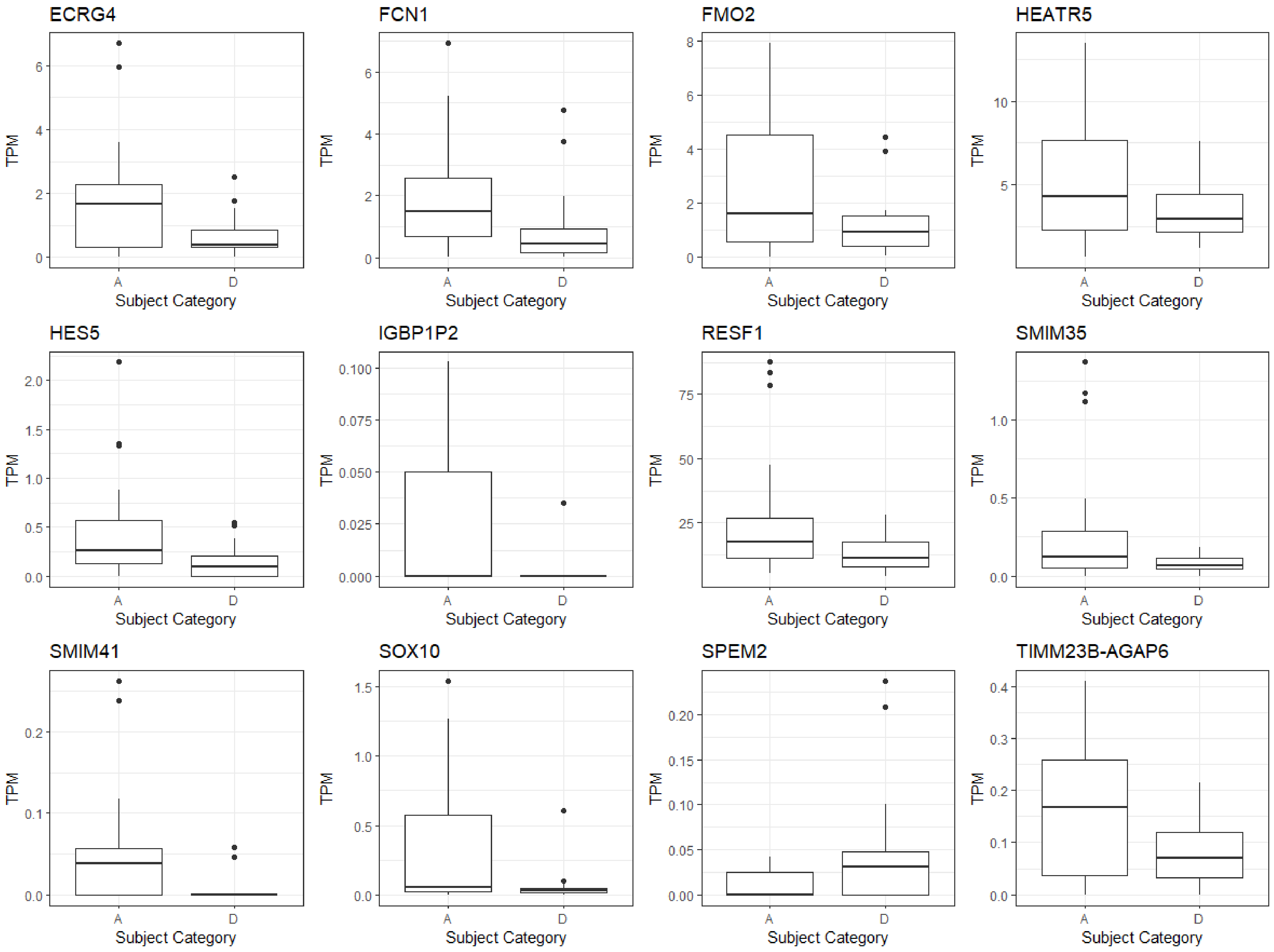

Supplement Figure 2C: Top Results of most significant diagnostic/prognostic biomarkers part 2

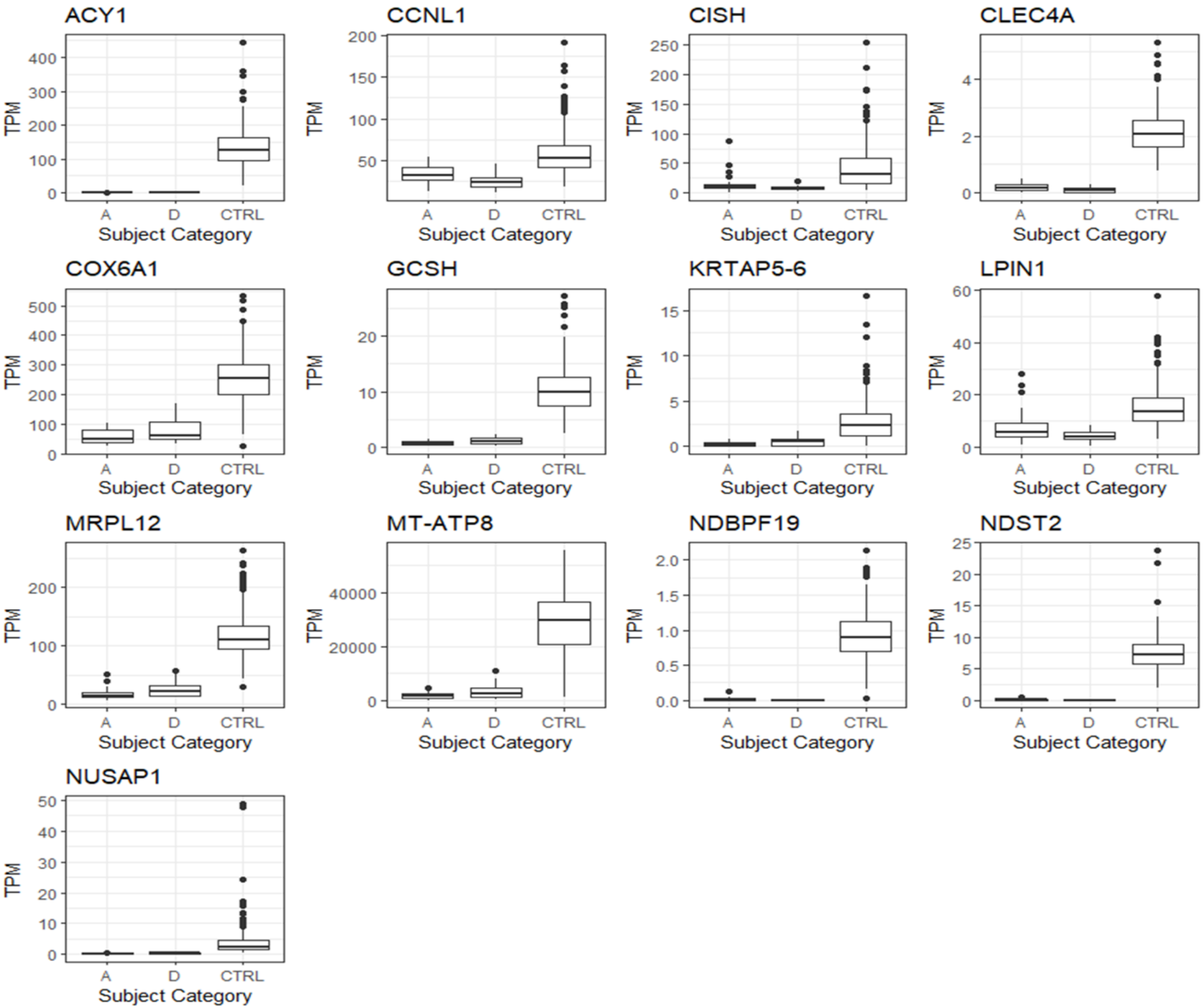

Supplemental Figure 3A: CCA Prognostic Gene Ontogeny

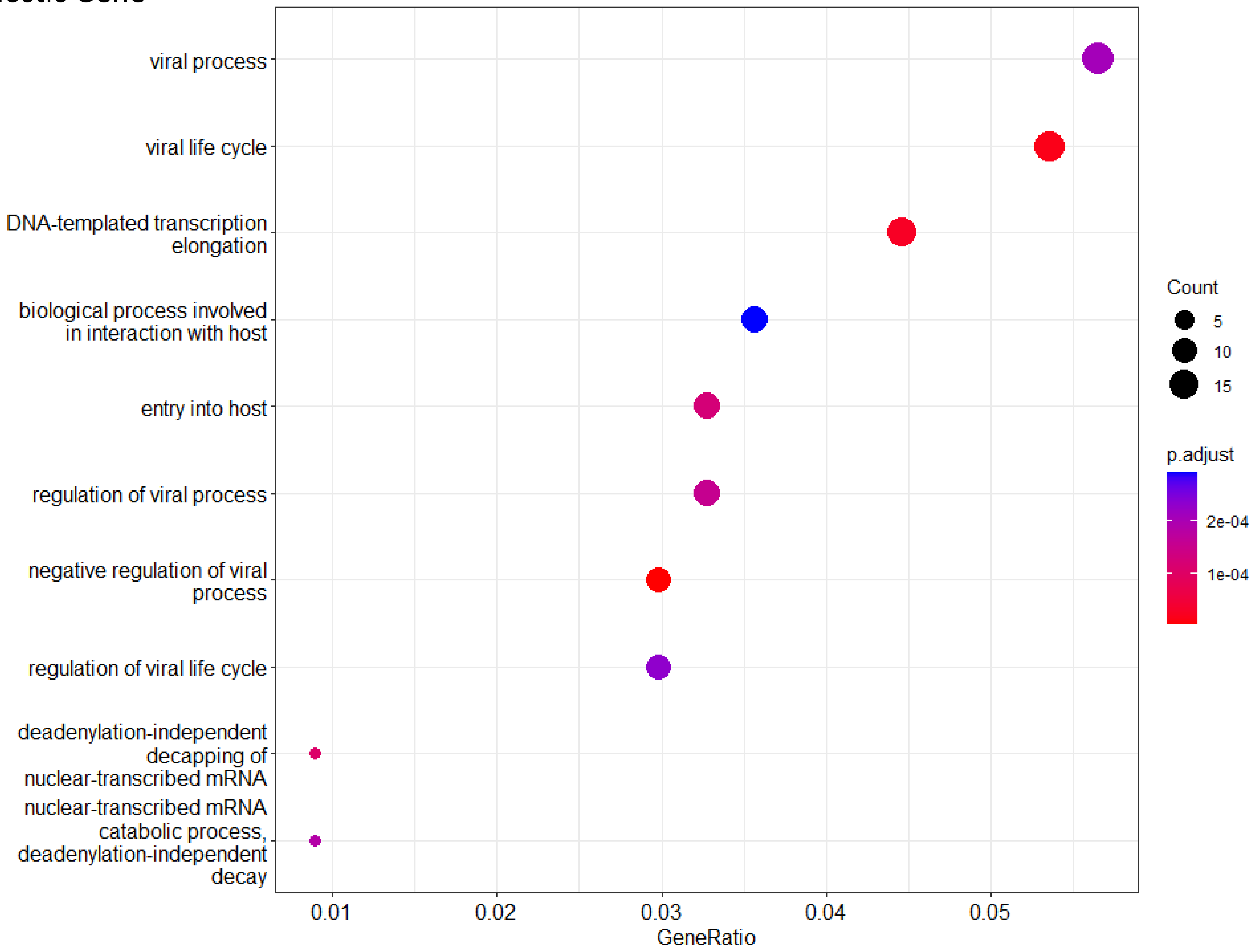

Supplemental Figure 3B: CCA Prognostic KEGG

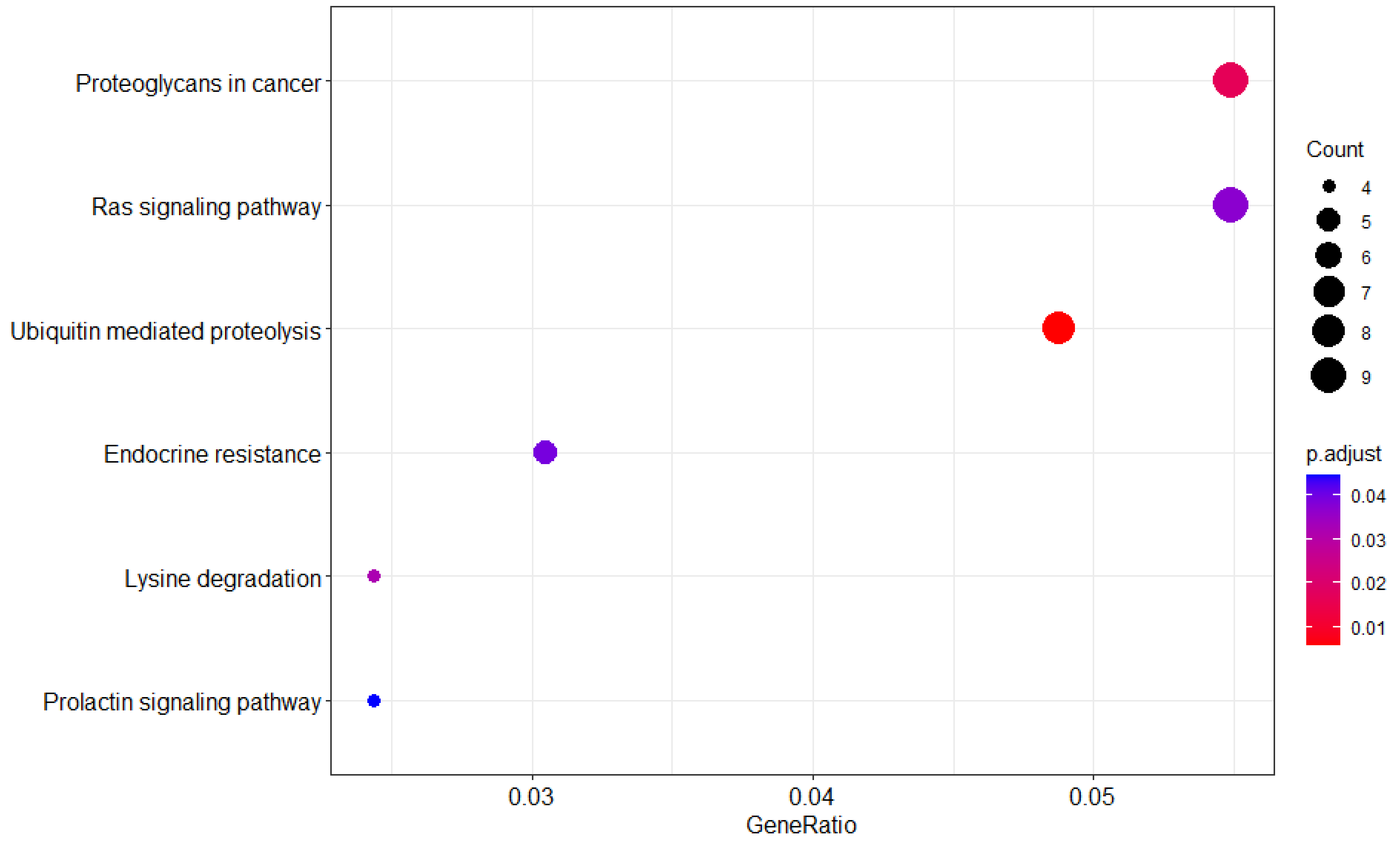

Supplemental Figure 3C: CCA Prognostic filter Disease Ontogeny

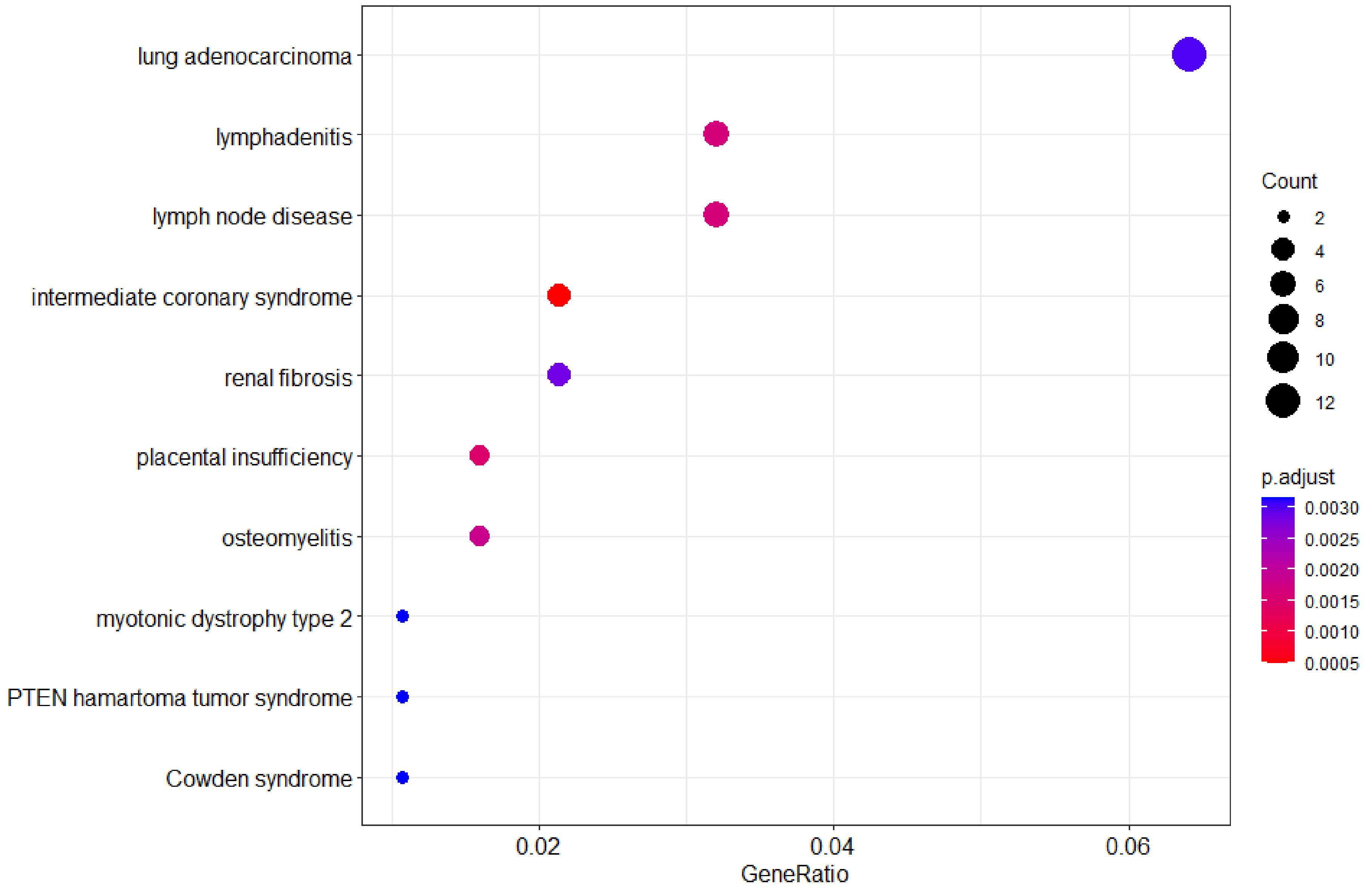

Supplemental Figure 4A: CCA Prognostic/Diagnostic filter Gene Ontogeny

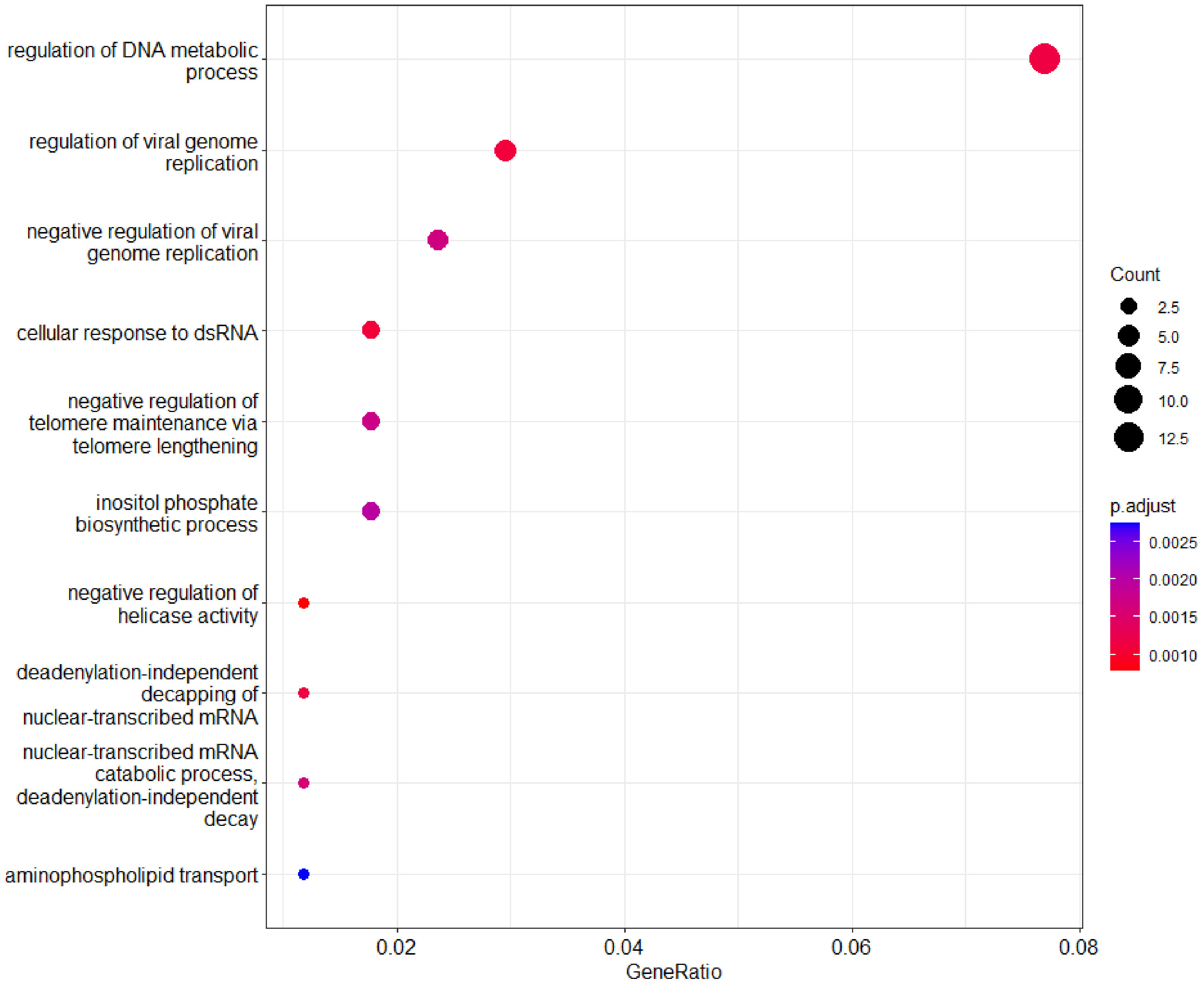

Supplemental Figure 4B: CCA Prognostic/Diagnostic filter KEGG

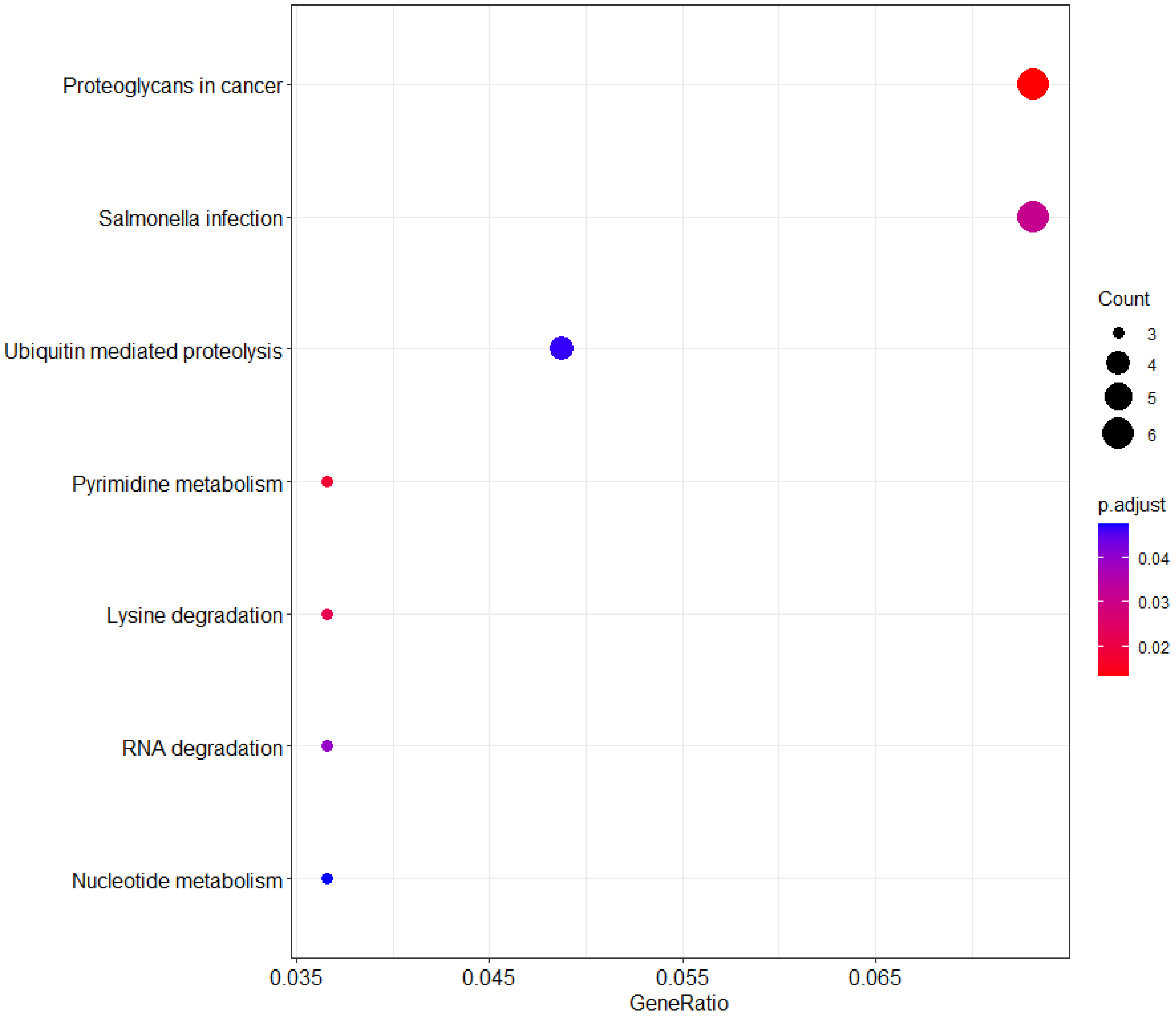

Supplemental Figure 4C: CCA Prognostic/Diagnostic filter Disease Ontogeny

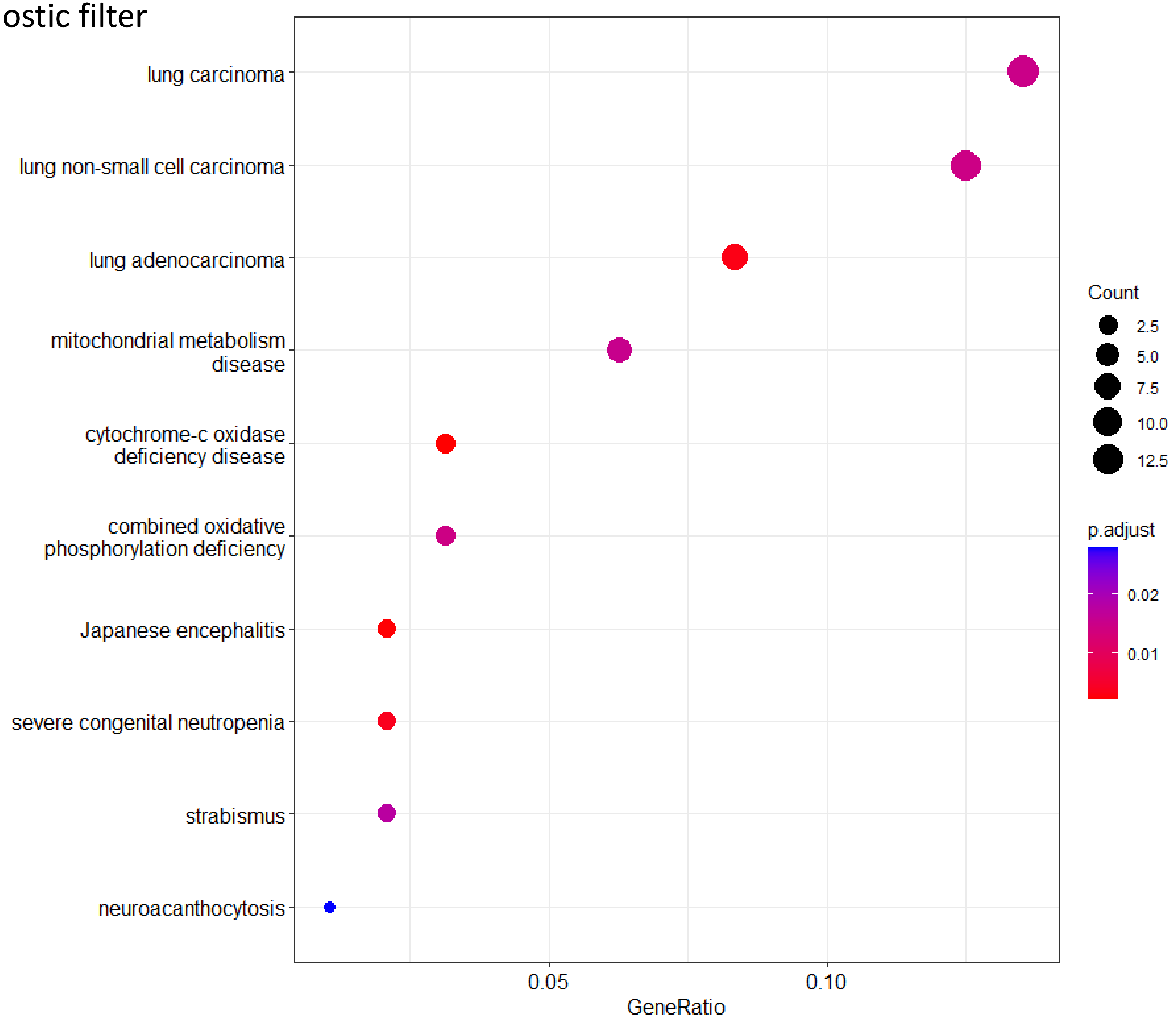

Supplemental Figure 5A: CCA predicted TF filter GSEA overexpression GO

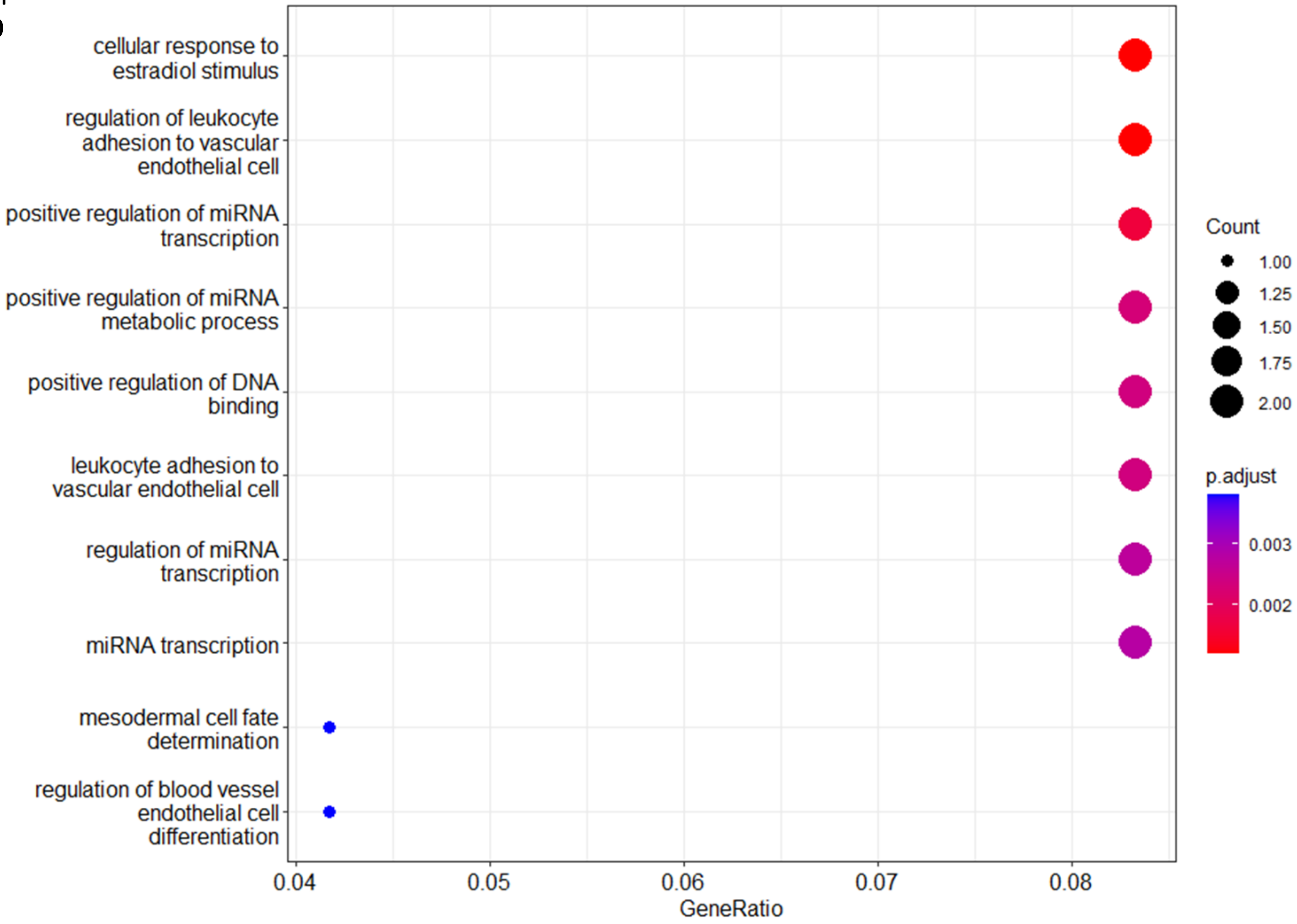

Supplemental Figure 5B: CCA predicted TF filter GSEA overexpression KEGG

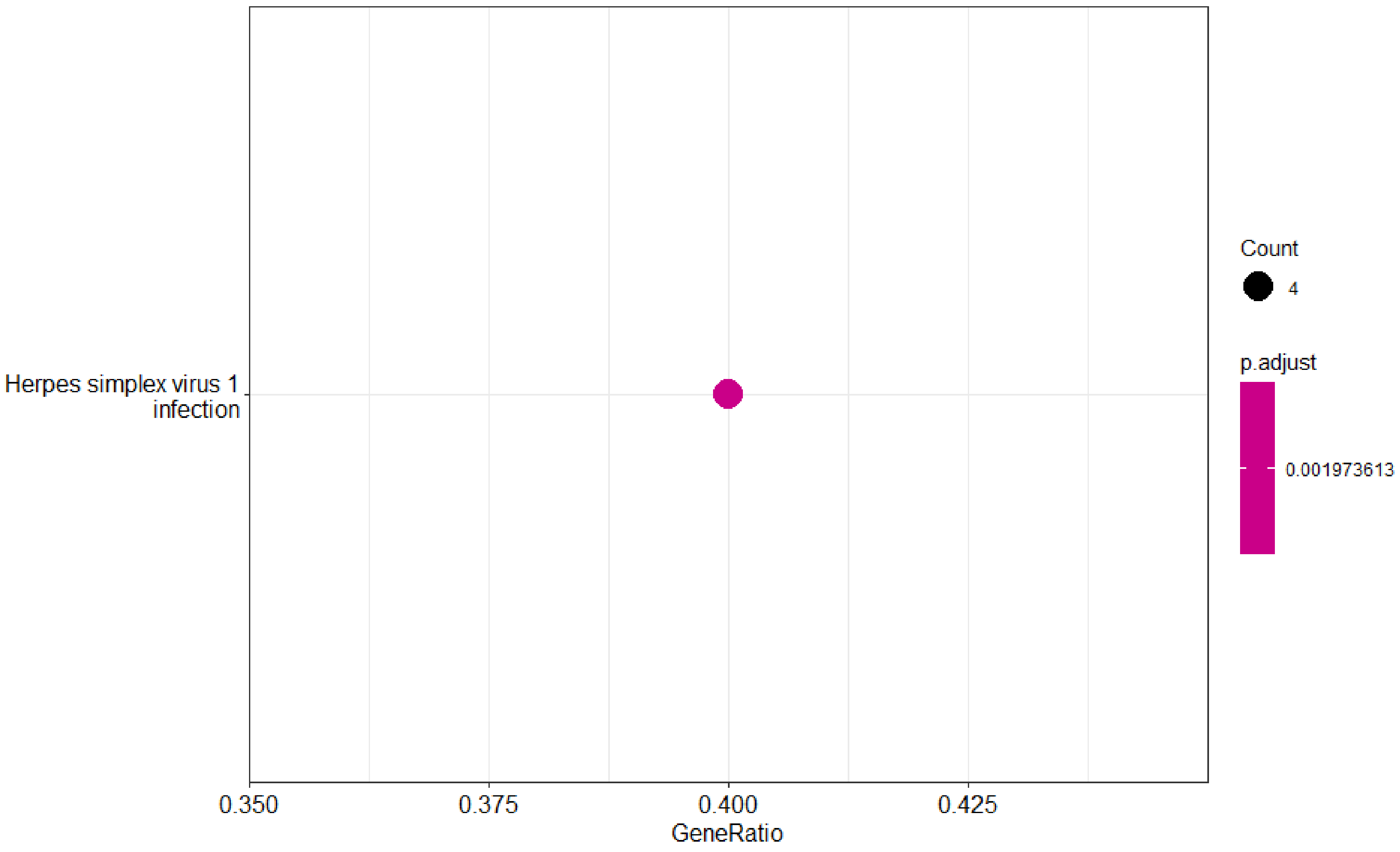

Supplemental Figure 5C : CCA Predicted TF filter GSEA overexpression Disease Ontogeny

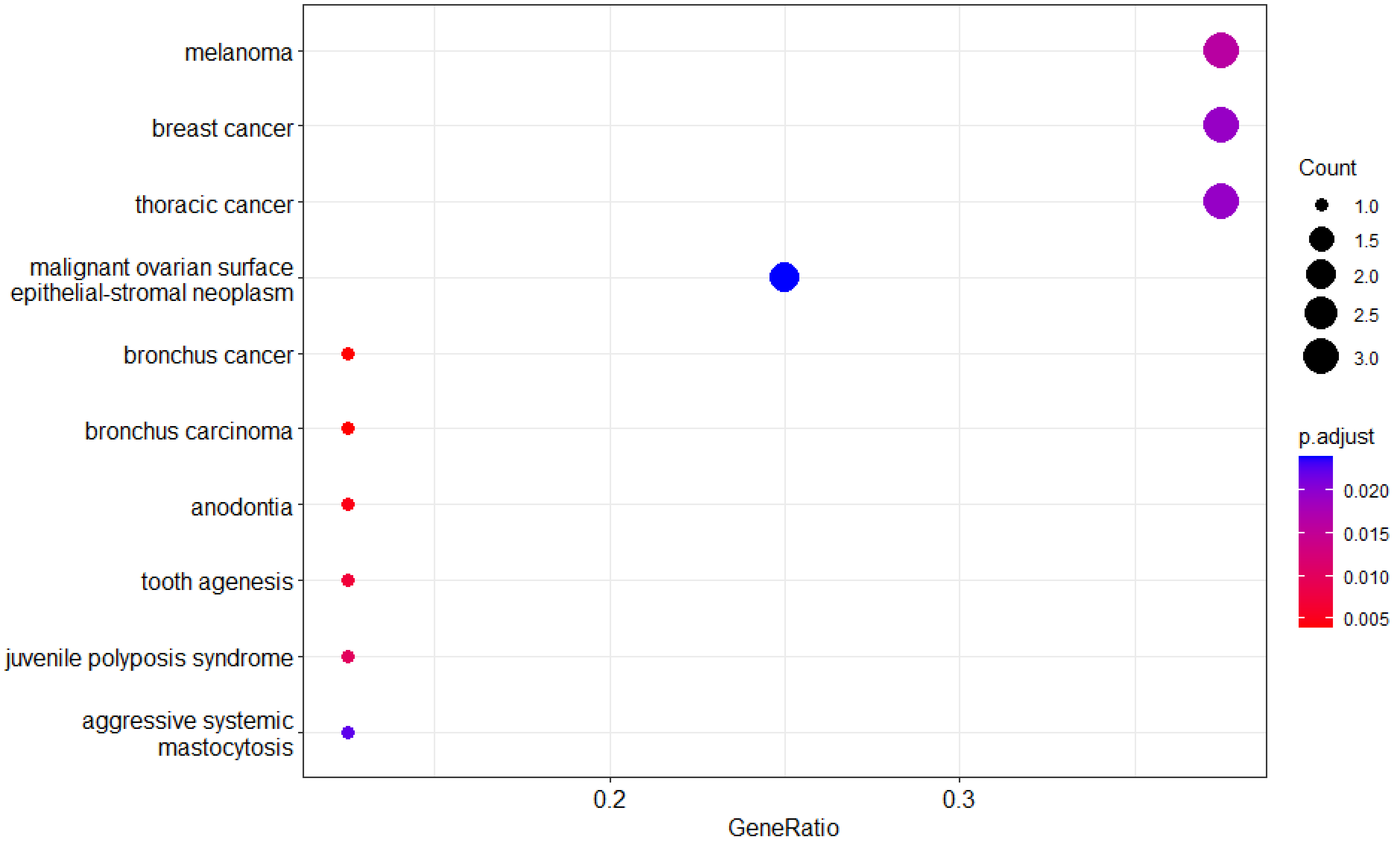
